# Epigenetic regulation of the circadian gene *Per1* in the hippocampus mediates age-related changes in memory and synaptic plasticity

**DOI:** 10.1101/301135

**Authors:** Janine L. Kwapis, Yasaman Alaghband, Enikö A. Kramár, Alberto J. López, Annie Vogel Ciernia, André O. White, Guanhua Shu, Diane Rhee, Christina M. Michael, Emilie Montellier, Yu Liu, Christophe N. Magnan, Paolo Sassone-Corsi, Pierre Baldi, Dina P. Matheos, Marcelo A. Wood

## Abstract

Aging is accompanied by impairments in both circadian rhythmicity and long-term memory. Although it is clear that memory performance is affected by circadian cycling, it is unknown whether age-related disruption of the circadian clock causes impaired hippocampal memory. Here, we show that the repressive histone deacetylase HDAC3 restricts long-term memory, synaptic plasticity, and learning-induced expression of the circadian gene *Per1* in the aging hippocampus without affecting rhythmic circadian activity patterns. We also demonstrate that hippocampal *Per1* is critical for long-term memory formation. Together, our data challenge the traditional idea that alterations in the core circadian clock drive circadian-related changes in memory formation and instead argue for a more autonomous role for circadian clock gene function in hippocampal cells to gate the likelihood of long-term memory formation.

## Background

Animals have an internal circadian clock that drives the rhythmic cycling of biological processes every ~24h. Circadian rhythms drive numerous physiological events, including the sleep-wake cycle, feeding behavior, body temperature, and metabolism. In the master circadian clock, the suprachiasmatic nucleus (SCN), a group of core clock genes oscillate in a negative feedback loop that cycles every ~24h (Deibel et al., 2015; Gerstner and Yin, 2010). In addition to regulating basic biological processes, the circadian clock also has a strong influence on memory. Long-term memory, which is transcription-dependent, shows a strong time-of-day effect, with peak memory performance during the day (inactive phase) in mice (Chaudhury and Colwell, 2002; Eckel-Mahan et al., 2008). Notably, both long-term memory and circadian rhythmicity are impaired with age (Kondratova and Kondratov, 2012), suggesting that these processes might share similar mechanisms. One idea is that clock genes located in memory-relevant structures, like the dorsal hippocampus, might gate an animal’s ability to form long-term memory based on the time of day (Rawashdeh et al., 2016). Consistent with this, disruption of several individual clock genes throughout the brain can impair hippocampal long-term memory in young animals. As no study to date has selectively disrupted circadian gene function within the dorsal hippocampus, it is unclear whether clock genes act within hippocampal cells to affect long-term memory formation or whether these memory deficits result from off-target effects in other brain regions, such as impaired circadian rhythms, sleep deficits, or even developmental abnormalities.

Gene expression is decreased in the aging brain, which may be the consequence of a more repressive chromatin structure. Transcription is controlled in part through changes in chromatin structure, which can dynamically promote or restrict access to neuronal DNA following a learning event. One hypothesis put forth by Barnes and Sweatt posits that the epigenome is altered in aging neurons, resulting in a repressive chromatin structure that prevents normal gene expression required for long-term memory formation (Penner et al., 2010). Several studies support this idea showing altered histone modification mechanisms in the aging brain (Peleg et al., 2010; Snigdha et al., 2016; Spiegel et al., 2014). However, whether chromatin modification mechanisms abnormally regulate circadian gene expression in a key learning and memory brain region is unknown. Here, we examined this possibility by focusing on the role of histone deacetylase 3 (HDAC3)-dependent regulation of age-related memory and gene expression.

## Results

### HDAC3 activity contributes to age-related hippocampal memory impairments

We first tested whether the repressive histone deacetylase HDAC3 plays a role in age-related memory decline. HDAC3 is a potent negative regulator of memory formation and disruption of HDAC3 in young animals can transform a subthreshold learning event into one generating persistent long-term memory for multiple tasks (Bieszczad et al., 2015; Kwapis et al., 2017b; Malvaez et al., 2013; McQuown et al., 2011). We used two methods of disrupting HDAC3 in the dorsal hippocampus of aging (18-month-old) mice. First, we created focal homozygous deletions of HDAC3 by infusing AAV2.1-CaMKII-Cre recombinase (1μL/side) into the dorsal hippocampi of HDAC3^flox/flox^ mice (Fig. S1A). Second, to selectively block the enzymatic activity of HDAC3, we used a dominant-negative point mutant virus, AAV2.1-CMV-HDAC3(Y298H)-v5 that specifically blocks HDAC3 deacetylase activity without affecting protein-protein interactions (Kwapis et al., 2017b; Lahm et al., 2007; Sun et al., 2013) Fig. S1B. Viruses were infused two weeks before training, allowing for tight spatial and temporal control over our HDAC3 manipulations to avoid potential side effects that might occur from prolonged HDAC3 disruption during development (Norwood et al., 2014; Nott et al., 2016). Two weeks after AAV-CaMKII-Cre infusion (Fig. S1C), we observed that *Hdac3* mRNA expression was not affected by training in object location memory (OLM), but genetic deletion of hippocampal *Hdac3* disrupted expression of *Hdac3* mRNA (Fig. S1D) in addition to HDAC3 protein (Fig. S1A).

To determine whether HDAC3 limits memory formation in the aging brain, we tested the effects of hippocampal HDAC3 deletion (HDAC3^flox/flox^) or activity disruption (HDAC3(Y298H)) on long-term memory for OLM (Fig. 1A). Consistent with numerous reports of age-related hippocampal memory deficits, we found that aging, 18-m.o. HDAC3^+/+^ mice displayed poor memory for OLM following 10-minute training (Fig. 1B, **gray bar**), which normally produces strong long-term memory in young mice (Vogel-Ciernia et al., 2013). In contrast, 18-m.o. HDAC3^flox/flox^ littermates formed robust long-term memory (Fig. 1B, **teal bar**), despite similar levels of total exploration (Fig. 1C). We observed similar effects with the activity-specific AAV-HDAC3(Y298H) virus. Aging, 18-m.o. empty vector (EV) control mice showed poor memory for OLM whereas mice infused with AAV-HDAC3(Y298H) into the DH showed significantly higher preference for the moved object with no change in total exploration (Fig. 1D to E). In contrast to the poor long-term memory observed in 18-m.o. wildtype mice (Fig. 1B and 1D), short-term memory for OLM (tested 60m after acquisition; Fig. S2A) was intact both groups of mice (Fig. S2B,C). We also observed no significant difference in movement or anxiety-like behavior between the groups (Fig. S2D-F). Together, these results demonstrate that age-related impairments in OLM are ameliorated by deletion or disruption of HDAC3 in the dorsal hippocampus.

**Fig. 1.**
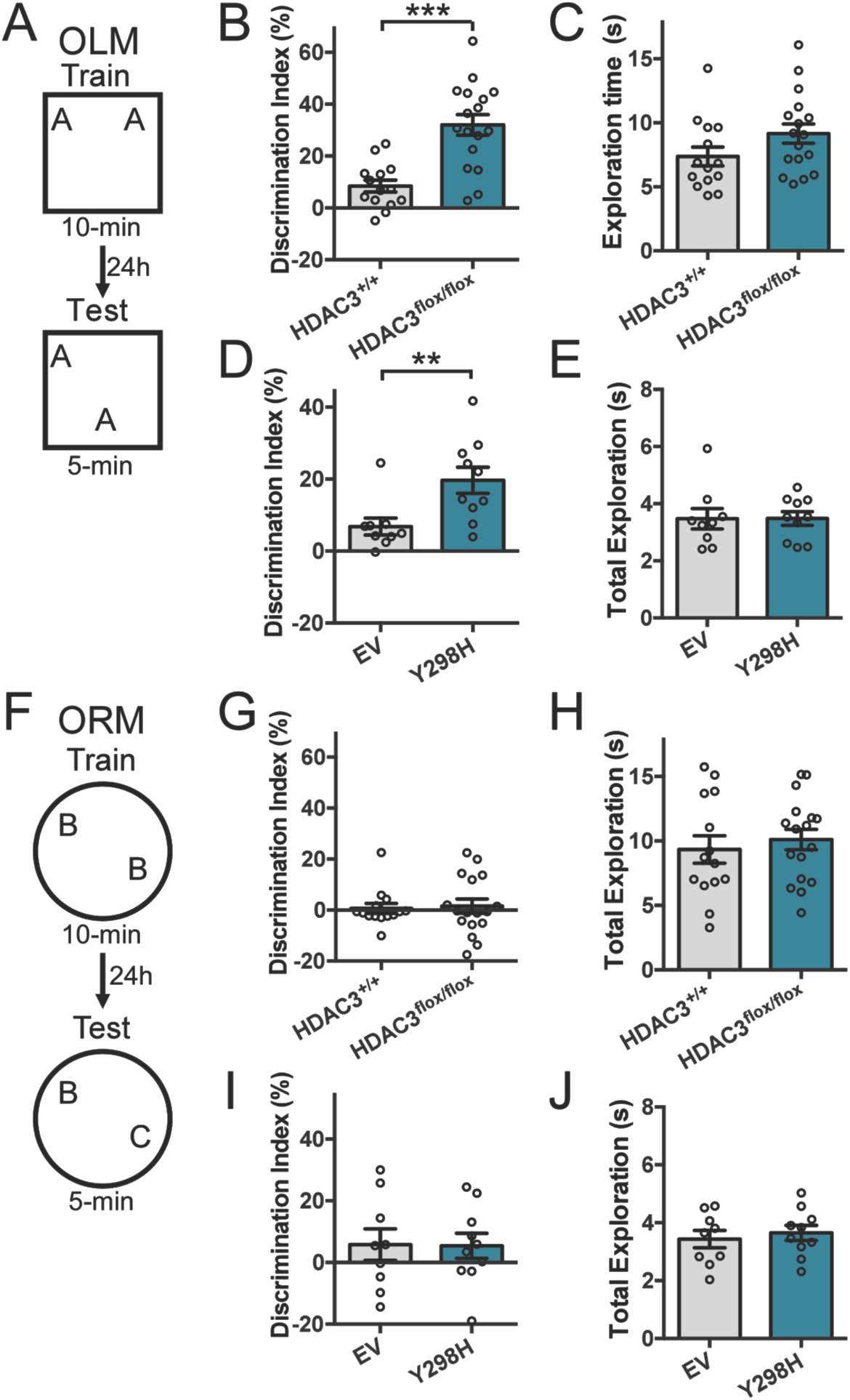
Deleting or disrupting HDAC3 ameliorates age-related impairments in hippocampal memory. **(A)** OLM procedure. AAV was infused 2 weeks before training. **(B)** 18-m.o. HDAC3flox/flox mice showed significantly better memory for OLM compared to HDAC3+/+ littermates (t(29) = 4.85, ***p<0.0001, n=14(5F), 17(6F)). **(C)** Total exploration was similar for both groups at test (t(29)=1.67, ***p=0.11). **(D)** Disrupting HDAC3 activity in the dorsal hippocampus with AAV-HDAC3(Y298H)-V5 also ameliorated hippocampal memory impairments in 18-m.o. mice (t(17)=2.9, **p<0.01, n=9,10; all males). **(E)** Total exploration time was similar for both groups at test (t(17)=0.02, p=0.98). **(F)** ORM experimental procedure, 2 weeks after the completion of OLM. **(G)** Both 18-m.o. HDAC3flox/flox mice and HDAC3+/+ littermates showed little preference for the novel object (t(29)=0.24, p=0.82, n=14(5F), 17(6F)). **(H)** Total exploration time was similar for both groups at test (t(29)=0.59, p=0.56). (I) Disrupting HDAC3 activity in the dorsal hippocampus with AAV-HDAC3(Y298H) also had no effect on ORM, with neither group showing preference for the novel object (t(17)=0.06, p=0.95, n=9,10; all males). **(J)** Groups showed similar total exploration time at test (t(17)=0.54, p=0.60). Data are presented as mean ±SEM.

We next tested whether our focal HDAC3 manipulation affected object recognition memory (ORM), which does not require the dorsal hippocampus for retrieval (Vogel-Ciernia et al., 2013). In this task, one familiar object is replaced by a novel item (Fig. 1F). Deleting HDAC3 in the dorsal hippocampus did not rescue memory for ORM, with both HDAC3^+/+^ and HDAC3^flox/flox^ mice showing no preference for the novel object at test (Fig. 1G). Again, groups did not differ in total exploration levels (Fig. 1H). Similarly, activity-specific disruption of HDAC3 in the hippocampus was unable to ameliorate age-related ORM impairments (Fig. 1I,J). Thus, age-related impairments in long-term ORM were not ameliorated by hippocampal deletion or disruption of HDAC3.

Together, our results indicate that deletion or disruption of HDAC3 in the dorsal hippocampus can ameliorate age-related long-term memory deficits for a hippocampus-dependent task (OLM; Fig. 1A-E) without affecting memory for a hippocampus-independent task (ORM; Fig. 2F-J). Importantly, all mice showed intact short-term memory for OLM (Fig. S2), suggesting that these animals acquire memory normally but fail to consolidate this information into observable long-term memory. As short-term memory is transcription-independent (for review, Alberini, 2009), this finding is consistent with the idea that learning-induced gene expression fails in aging mice, resulting in age-related impairments in long-term memory. Together, these findings are consistent with the hypothesis that HDAC3 contributes to a repressive chromatin structure in the aging hippocampus that limits memory formation (Penner et al., 2010).

**Figure 2.**
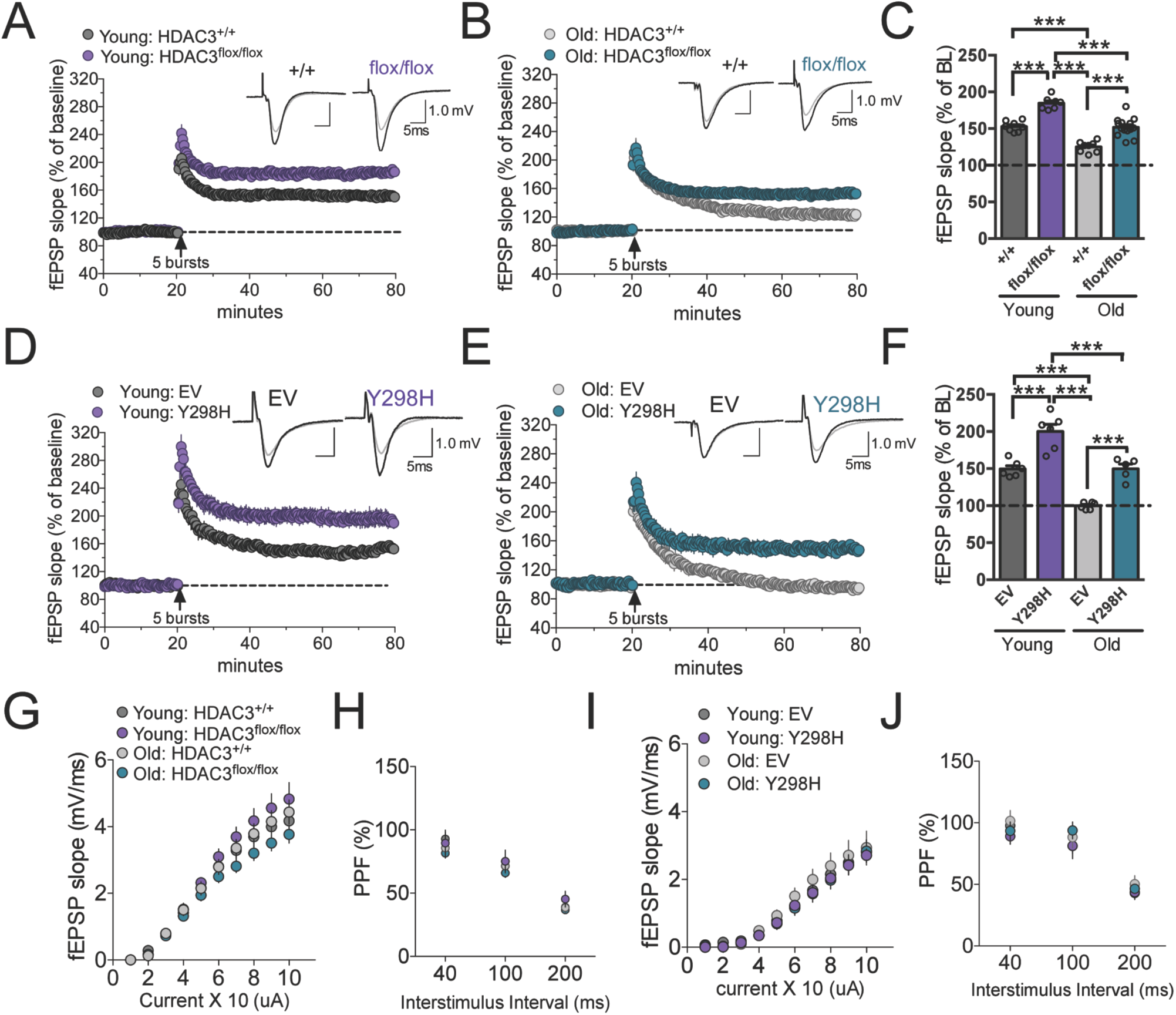
Deleting or disrupting HDAC3 ameliorates age-related impairments in synaptic plasticity. **(A)**Mean±SEM fEPSP slope recordings in hippocampal slices from young (3-m.o.) HDAC3flox/flox mice or HDAC3+/+ littermates 2-weeks after hippocampal AAV-Cre infusion. Deleting HDAC3 enhanced theta burst-induced LTP in the young hippocampus. **(B)** The same stimulation caused a gradual decay toward baseline in slices from old (18-m.o.) HDAC3+/+ mice. Deleting HDAC3 (HDAC3flox/flox) restored a greater level of stable potentiation relative to HDAC3+/+ littermates. **(C)** Summary graph showing mean fEPSP slope 60m after stimulation. Potentiation was significantly enhanced by HDAC3 deletion in slices from both young and old mice. Deleting HDAC3 in the 18-m.o. hippocampus produced LTP comparable to that of 3-m.o. wildtype mice (Two-way ANOVA: main effects of Age (F(1,33)=80.8, p<0.0001) and Genotype (F(1,33)=75.5, p<0.0001), Sidak’s post hoc tests, ***p < 0.0001, n=8,7,8,14 slices from 3,3,3,6 mice; all male). **(D)** Mean±SEM fEPSP slope recordings in hippocampal slices from young (3-m.o.) mice two weeks after hippocampal AAV-HDAC3(Y298H) or AAV-EV infusion. Disrupting HDAC3 activity enhanced LTP in the young hippocampus. **(E)** The same stimulation protocol failed to produce stable potentiation in slices from 18-m.o. mice, but this impairment was overcome by disrupting HDAC3 activity (AAV-HDAC3(Y298H)). **(F)** Summary graph. Potentiation was significantly enhanced by HDAC3 disruption in slices from both young and old mice. Disrupting HDAC3 activity in the 18-m.o. hippocampus with AAV-HDAC3(Y298H) produced LTP comparable to that of 3-m.o. wildtype mice (Two-way ANOVA: main effects of Age (F(1,19)=62.8, p<0.0001) and Genotype (F(1,19)=63.4, p<0.0001), Sidak’s post hoc tests, ***p<0.0001, n=6,6,6,5 slices from 4,4,4,4 mice; all male). **(G)** Input/output (I/O) curves and **(H)** paired-pulse facilitation (PPF) were comparable between HDAC3+/+ and HDAC3-flox/flox slices. **(I)** I/O curves and **(J)** PPF were similar between AAV-EV and AAV-HDAC3(Y298H) slices. Data are presented as mean ±SEM.

### Deletion or disruption of HDAC3 also ameliorates age-related deficits in LTP

To test whether HDAC3 also contributes to age-related synaptic plasticity impairments, we examined long-term potentiation (LTP) in acute hippocampal slices following either deletion or disruption of HDAC3. LTP is also impaired with age, particularly when the stimulation protocol is close to the LTP induction threshold (Barnes, 2003). Two weeks after viral infusion, we prepared hippocampal slices and induced LTP with a single train of 5 theta bursts to Schaffer collateral inputs and recorded field EPSPs from apical dendrites of CA1b. This relatively mild form of stimulation produced a stable level of LTP in young HDAC3^+/+^ slices (Fig. 2A).

Deleting HDAC3 in the hippocampus enhanced LTP, with HDAC3^floxflox^ mice showing significantly higher potentiation than wildtype controls. As predicted, aging-18-m.o. HDAC3^+/+^ mice showed impaired LTP and the HDAC3 deletion ameliorated this deficit, producing LTP comparable to that of young wildtype mice (Fig. 2B,C) with no effect on baseline synaptic transmission (Fig. 2G,H).

We observed similar results with the activity-specific disruption. Here, we used a within-subjects design in which young and old wildtype mice were infused with the control virus (AAV-EV) into one hippocampus and AAV-HDAC3(Y298H) into the contralateral hippocampus. As before, we found that disrupting HDAC3 activity enhanced LTP in young mice (Fig. 2D) and ameliorated age-related LTP impairments in aging mice (Fig. 2E,F) without interfering with baseline synaptic transmission (Fig. 2I-J). Either deletion or disruption of HDAC3 can therefore ameliorate age-related impairments in hippocampal LTP.

### Age-related changes in gene expression are partially rescued through HDAC3 deletion

Our data suggest that deleting or disrupting HDAC3 ameliorates age-related impairments in long-term memory and synaptic plasticity. We next asked whether age-related deficits in hippocampal gene expression could also be ameliorated by deleting HDAC3. We hypothesized that dysregulation of HDAC3 activity in the old brain leads to an unusually repressive chromatin structure that limits gene expression, which ultimately impairs long-term memory. To identify which specific genes are regulated by HDAC3 in the young and aging brain, we ran an RNA sequencing experiment in which we compared young (3-m.o.) wildtype mice, aging (18-m.o.) HDAC3^+/+^ mice, and aging (18-m.o.) HDAC3^flox/flox^ littermates (Fig. 3A). To identify gene expression changes during memory consolidation, animals in each group were killed 60m after 10-min OLM training and compared to homecage (HC) controls. After mapping and considering the haploid genome (Langmead et al., 2009; Speir et al., 2016), sequencing quality was assessed (Fig. S3A,B) and significant differences in expression profiles were examined between all pairs of samples for *p* < 0.05 (Vogel-Ciernia et al., 2013).

**Figure 3.**
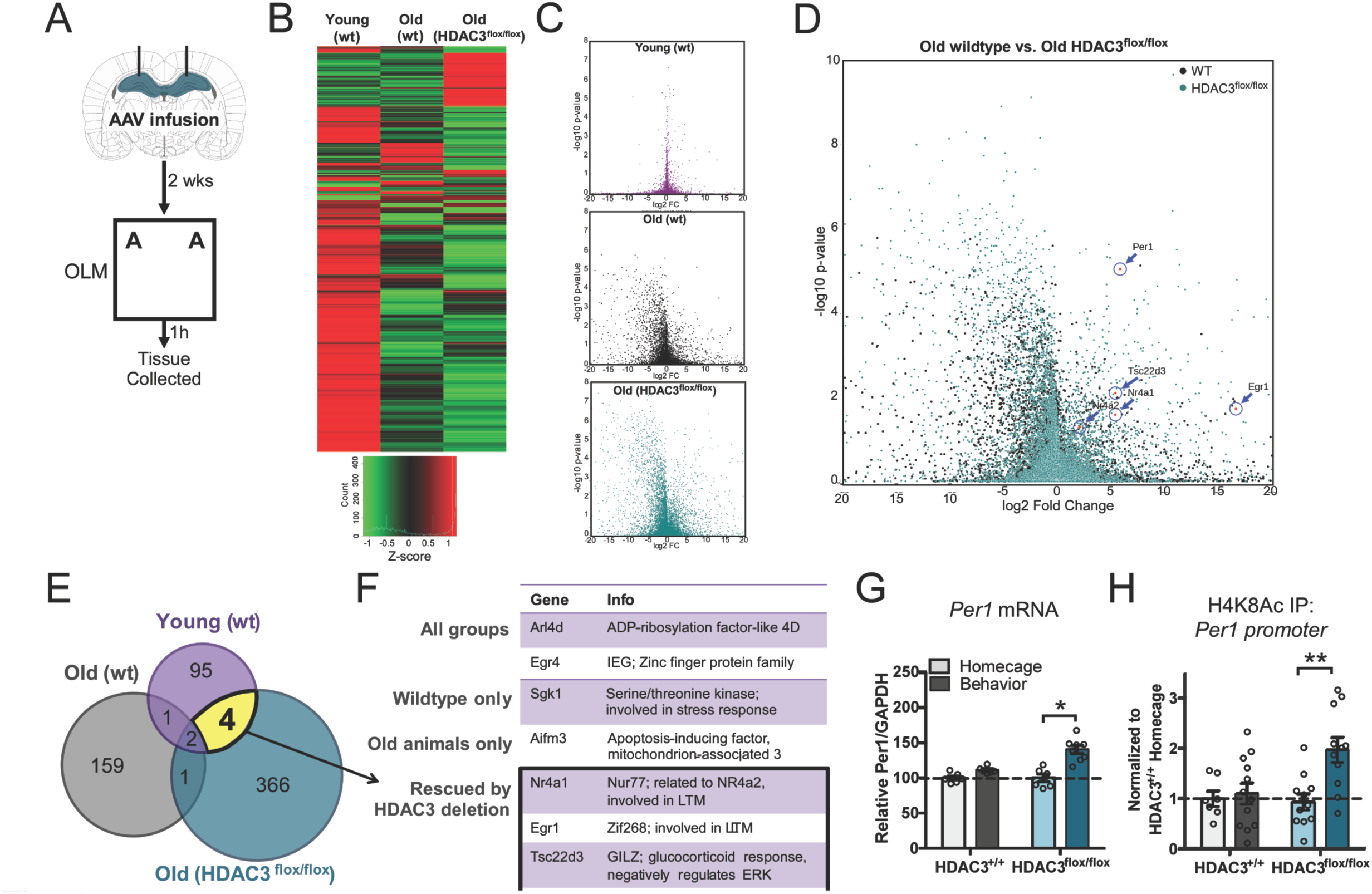
A subset of learning-induced genes are impaired with age and rescued by HDAC3 deletion. **(A)** Experimental procedure. **(B)** Heat map comparing learning-induced gene changes in each group. Each row represents an individual gene. Color legend shown at bottom; red, upregulation; green, downregulation. **(C)** Volcano plots illustrating the significance (Y-axis) and magnitude (X-axis) of learning-induced changes in each group. Young animals showed fewer learning-induced changes in gene expression compared to old wildtype or old HDAC3flox/flox mice. **(D)** Overlapping volcano plot comparing old wildtype and old HDAC3flox/flox groups. Per1 was strongly induced by learning in HDAC3flox/flox mice. **(E)** Gene expression diagram displaying genes expressed higher in trained animals compared to homecage controls. Total gene count shown inside circles. **(F)** Table listing learning-induced genes common to two or more groups. Of particular interest are the four genes “rescued by HDAC3 deletion” that are upregulated in the young wildtype and old HDAC3flox/flox groups but are not upregulated in the old wildtype group. **(G)** Per1 mRNA expression is increased in the 18-m.o. brain in the absence of HDAC3 (Genotype × Train, F(1,22)=9.48, *p<0.05; n=6(2F),6(3F),6(4F),8(5F)). **(H)** Occupancy of H4K8Ac at the Per1 promoter is upregulated by learning in the 18-m.o. hippocampus in the absence of HDAC3 (Genotype × Train, F(1,37)=5.1, **p=0.01, n=7(4F),12(4F),11(4F),11(6F)). Data are presented as mean ±SEM.

We expected that learning-induced gene expression would be altered in the old brain, as previously described, with a subset of genes failing to express after learning. We therefore focused on those genes expressed at significantly higher levels in the trained groups compared to homecage controls. While each group (Young WT, Old WT, Old HDAC3^flox/flox^) showed a substantial number of genes induced by learning, each group showed a unique gene expression profile (Fig. 3B, Tables S1–3). Old brains (both wildtype and HDAC3^flox/flox^) showed a greater number of up- and down-regulated genes in response to learning than young wildtype brains (Fig. 3C), with a bias towards general gene repression (Fig. 3B) as previously observed (Rowe et al., 2007), suggesting that learning-induced gene expression is less tightly regulated in the aging brain. Most of the genes upregulated by learning were unique to that group, with little overlap between the three conditions (Fig. 3D, 3E). This indicates that the aging brain is characterized by both failed induction of genes that are typically upregulated by learning *and* aberrant induction of genes that are not usually expressed during memory consolidation. Further, when HDAC3 is deleted from the aging brain, learning induces a unique gene expression profile, rather than recapitulating the gene expression profile of the young brain.

Only a handful of genes were common to two or more groups (Fig. 3E,F). Of particular interest is the subset of genes upregulated in both the young wildtype and old HDAC3^flox/flox^ groups that do not show learning-induced increases in the old wildtype hippocampus. These are the genes that fail to normally express in the old brain after learning but are “rescued” by HDAC3 deletion and may therefore function as a mechanism through which HDAC3 deletion ameliorates age-related memory impairments. Four genes were identified in this group: *Nr4a1*, *Egr1*, *Tsc22d3*, and *Per1*. All of these genes have been broadly implicated in memory formation (McNulty et al., 2012; Rawashdeh et al., 2016; Vecsey et al., 2015), although this is the first study to demonstrate that learning-induced expression of these genes is impaired with age and rescued with HDAC3 deletion.

### Deleting HDAC3 ameliorates deficits in *Per1* expression and acetylation at the *Per1* promoter without affecting circadian rhythmicity

Of the genes identified through our RNA-seq, *Period1* (*Per1*) was the most strongly induced by learning in the HDAC3^flox/flox^ group (Fig. 3D). This target is particularly intriguing, as *Per1* is typically studied in the context of circadian rhythms but has also been implicated in hippocampal memory formation (Jilg et al., 2010; Rawashdeh et al., 2014; Rawashdeh et al., 2016). As aging is known to be accompanied by impairments in circadian rhythms (Kondratova and Kondratov, 2012) and memory is linked to time-of-day (Eckel-Mahan, 2012; Gerstner and Yin, 2010), this target of HDAC3 may represent a critical and underexplored interface between aging, chromatin modification, and the circadian clock.

To further examine the expression of *Per1* in old HDAC3^+/+^ and old HDAC3^flox/flox^ animals, we used RT-qPCR and ChIP-qPCR. We found that learning failed to induce upregulation of *Per1* in the dorsal hippocampus of 18-m.o. HDAC3^+/+^ mice, but in the absence of HDAC3 (HDAC3^flox/flox^), learning triggered a significant increase in *Per1* mRNA (Fig. 3G). We also measured the expression of two additional genes that play a well-documented role in long-term memory formation: *Arc* and *cFos* (Guzowski et al., 2001). Learning-induced expression of *Arc* was intact in the aging wildtype brain and was unaffected by HDAC3 deletion (Fig. S3C). *cFos* expression, on the other hand, failed to be induced by learning in the old HDAC3^+/+^ hippocampus, but deleting HDAC3 was not sufficient to restore this failed induction (Fig. S3D). Deleting HDAC3 therefore only restores expression of a subset of learning-induced genes in the aging brain, including *Per1.*

To determine whether deleting HDAC3 restores expression of *Per1* by promoting histone acetylation along its promoter, we next measured acetylation of histone 4, lysine 8 (H4K8Ac) at *Per1* CRE promoter site using chromatin immunoprecipitation (ChIP-qPCR). H4K8Ac is a marker of transcriptional activation (Kouzarides, 2007) and is also a known target of HDAC3 (Kwapis et al., 2017b; Malvaez et al., 2013). Learning did not change H4K8Ac levels at the *Per1* promoter in the old wildtype brain but in the absence of HDAC3, H4K8Ac levels at the *Per1* promoter were significantly increased in response to learning (Fig. 3H). For *Arc* and *cFos*, we saw no change H4K8Ac occupancy across groups (Fig. S3E,F). Together, these results suggest that deleting HDAC3 restores acetylation at the *Per1* promoter and expression of *Per1* mRNA in response to learning.

One outstanding question is whether the observed changes in *Per1* are due to changes in the circadian rhythm of HDAC3^flox/flox^ mice. If deletion of HDAC3 in the dorsal hippocampus alters circadian rhythmicity, this could explain the observed changes in both *Per1* expression and long-term memory formation. To rule this out, we assessed the circadian rhythmicity of young (3-m.o.) and aging (18-m.o.) HDAC3^flox/flox^ mice and their HDAC3^+/+^ littermates following AAV-CaMKII-Cre infusion. After 2 weeks of entrainment to a 12h light/dark cycle (LD), mice were put in constant darkness (DD) to measure endogenous circadian rhythms (Fig. S4A). We observed no difference in circadian activity patterns between HDAC3^+/+^ and HDAC3^flox/flox^ mice at either age group (Fig. S4B to S4F), suggesting that hippocampal HDAC3 has no effect on circadian rhythmicity. Thus, the observed changes in *Per1* expression and long-term memory formation following HDAC3 deletion cannot be explained by changes in circadian rhythmicity.

### *Per1* is induced by learning and regulated by HDAC3

To test whether hippocampal *Per1* is required for long-term memory formation, we assessed whether hippocampus-dependent learning typically induces *Per1* mRNA expression. We sacrificed young (3-m.o.) wildtype mice 60m after acquisition of either OLM or context fear conditioning (CFC) (Kwapis et al., 2017b). *Per1* mRNA expression was significantly upregulated in animals trained with either OLM (Fig. 4A) or CFC (Fig. 4B) compared to homecage controls, indicating that *Per1* mRNA expression is typically induced in the hippocampus during memory consolidation for multiple tasks.

**Figure 4.**
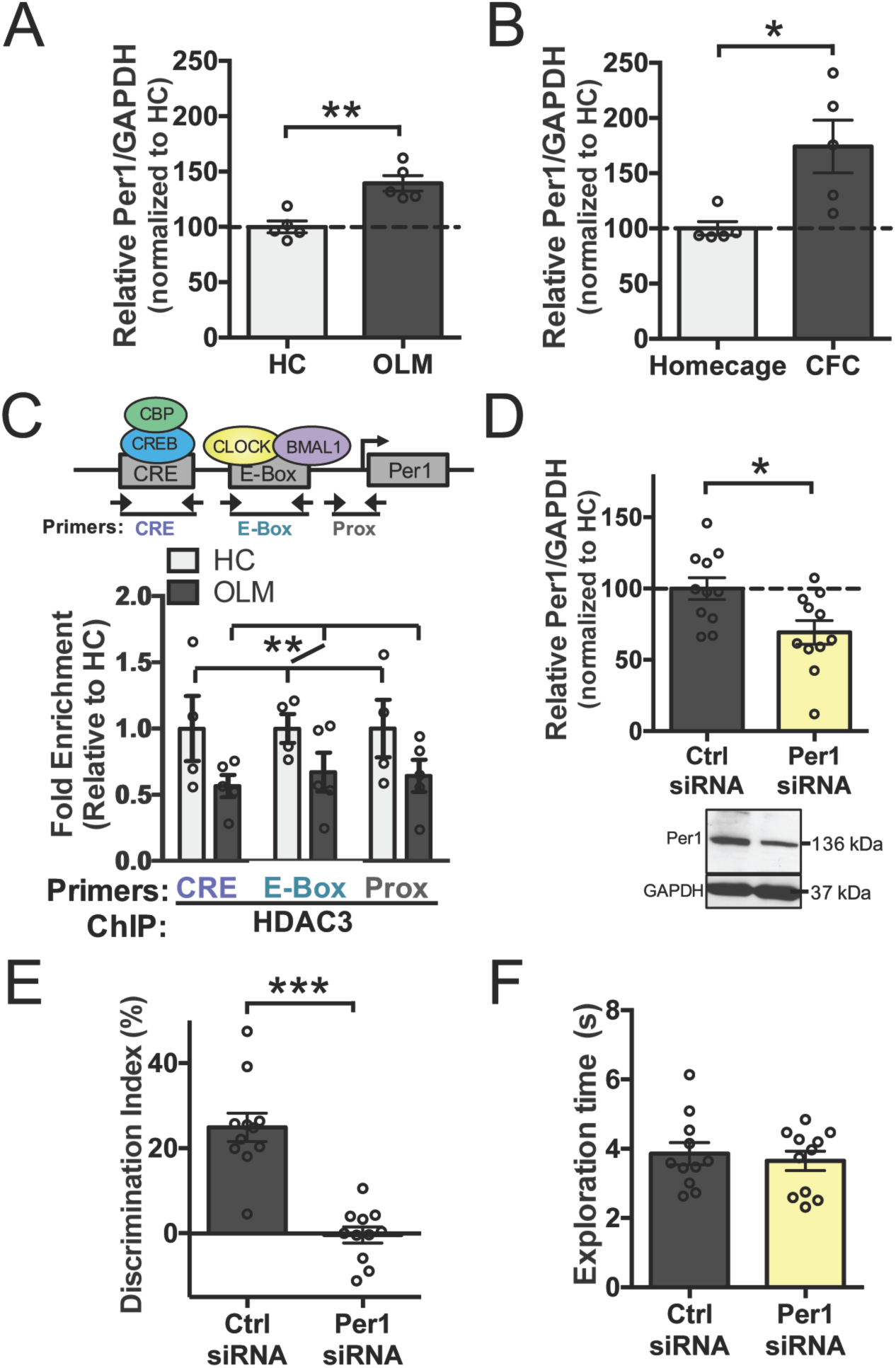
Knockdown of PER1 in the dorsal hippocampus impairs long-term memory. Per1 mRNA expression is upregulated 60m in the young hippocampus after **(A)** OLM (t(8)=4.49, **p<0.01; n=5,5; all male) and **(B)** context fear conditioning (t(8)=3.01, *p<0.05, n=5,5; all male). **(C)** Top, schematic of ChIP primer sites along the Per1 promoter. Bottom, H4K8Ac ChIP results. HDAC3 occupancy was reduced along the Per1 promoter in the young hippocampus after OLM (Training effect only, F(1,21)=8.51, **p<0.01, n=4,5; all male). **(D)** Hippocampal PER1 protein expression was significantly reduced by Per1 siRNA infusion (t(20)=2.72, p=0.01, n=11,11; all male). **(E)** Per1 knockdown impaired OLM (t(20)=6.58, ***p>0.0001, n=11,11; all male). **(F)** Per1 siRNA did not affect total exploration time at test (t(20)=0.049, p=0.63). Data are presented as mean ±SEM.

To determine whether this learning-induced increase in *Per1* might be mediated through HDAC3, we next used ChIP-qPCR to measure HDAC3 occupancy after learning at different sites along the *Per1* promoter in the young hippocampus (Fig. 4C, **top**). We found that HDAC3 occupancy at the *Per1* promoter was reduced at all three tested sites following OLM training (Fig. 4C, **bottom**). Along with our previous finding that HDAC3 deletion restores both acetylation at *Per1* and *Per1* mRNA expression (Fig. 3), this strongly suggests that HDAC3 regulates *Per1* expression in the dorsal hippocampus and dysregulation of HDAC3 may contribute to age-related impairments in learning-induced *Per1* expression. PER1 may therefore be part of a key mechanism through which HDAC3 deletion ameliorates age-related impairments in hippocampal long-term memory formation.

### Knockdown of *Per1* impairs long-term memory in young mice

Next, to test whether learning-induced upregulation of *Per1* in the dorsal hippocampus is necessary for long-term memory formation, we infused siRNA targeting *Per1* into the dorsal hippocampus of young mice 48h before 10-minute OLM training. Infusion of *Per1* siRNA produced a significant reduction of PER1 protein in the dorsal hippocampus, as measured 2h after the final test session (Fig. 4D). This relatively modest knockdown of PER1 protein (~30%) produced severely impaired memory formation for OLM; mice infused with *Per1* siRNA showed significantly less preference for the moved object compared to control mice (Fig. 4E) with no effect on total object exploration (Fig. 4F). This demonstrates, for the first time, that local disruption of a core circadian clock gene selectively within the hippocampus can impair memory formation.

### Overexpression of *Per1* ameliorates age-related hippocampal memory impairments

Finally, to determine whether overexpression of *Per1* in the dorsal hippocampus is sufficient to ameliorate age-related memory impairments, we locally upregulated *Per1* using two complementary methods. First, we used a lentivirus expressing wildtype *Per1* with a v5 epitope tag (pLVX-v5Per1) (Fig. 5A). Infusion of pLVX-v5Per1 into the dorsal hippocampus of 18-m.o. mice 2 weeks before behavior significantly improved performance on the OLM task relative to pLVX-EV (empty vector) controls (Fig. 5B) without affecting total exploration (Fig. 5C).

**Figure 5.**
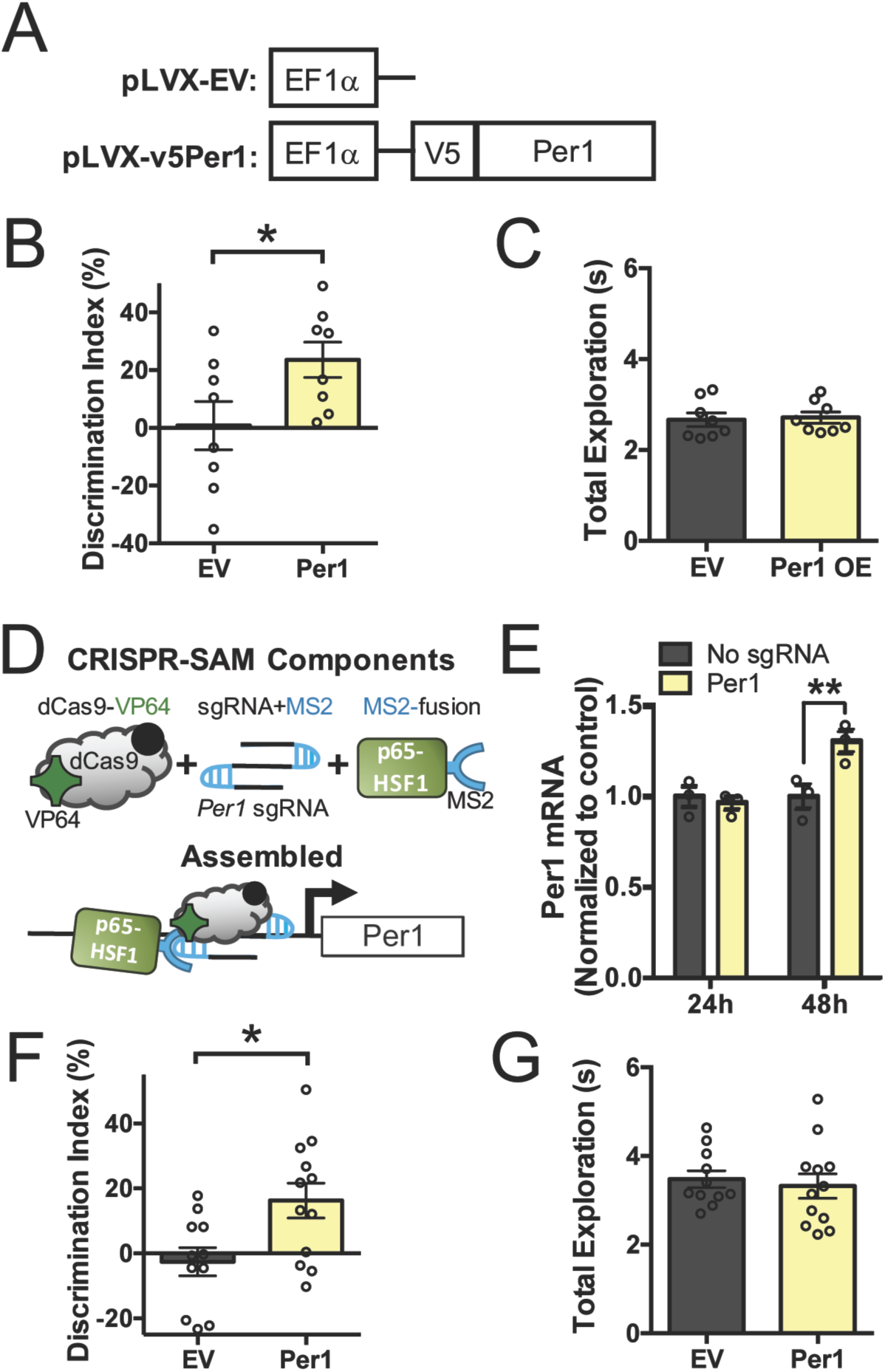
Overexpression of Per1 in the dorsal hippocampus ameliorates age-related impairments in object location memory. **(A)** Schematic of lentivirus construct used to overexpress v5-tagged Per1 (pLVX-v5Per1) compared to the empty vector control (pLVX-EV). **(B)** 18-m.o. mice given hippocampal Infusions of pLVX-v5Per1 showed significantly better memory for OLM than mice given pLVX-EV control virus (t(14) = 2.2, *p<0.05, n=8,8, all males). **(C)** Total exploration was similar for both groups at test (t(14) = 0.24, p=0.8). **(D)** Schematic of CRISPR/dCas9 Synergistic Activation Mediator (SAM) system used to drive Per1 transcription. Top: Individual components of SAM. Bottom: Components assembled at the Per1 promoter, driving Per1 transcription. **(E)** Per1 mRNA was significantly increased 48h after transfection of CRISPR-SAM components in HT22 cells compared to cells transfected without sgRNA (Two-way ANOVA: Group × Timepoint interaction, (F(1,8)=8.98, p<0.05), Sidak’s post hoc tests, *p<0.05, n=3,3,3,3). **(F)** 18-m.o. mice given hippocampal infusions of the CRISPR-SAM system with sgRNA targeting Per1 showed significantly better memory for OLM compared to EV control mice without sgRNA (t(21) = 2.7, *p<0.05). **(G)** Total exploration was similar for both groups at test (t(21) = 0.45, p=0.66). Data are presented as mean ±SEM.

To complement this approach, we also used the CRISPR/dCas9 Synergistic Activation Mediator (SAM) system (Konermann et al., 2015) to drive transcriptional activation of *Per1* in the dorsal hippocampus. This system consists of three lentiviral components: a catalytically inactive Cas9 (dCas9) fused to a VP64 transcriptional activation domain, a modified single-guide RNA (sgRNA) targeting *Per1* with two MS2 RNA adaptamers that can recruit the third component, an MS2-double transcriptional activator (MS2-p65-HSF1) fusion protein (Fig. 5D). Control animals received an empty vector in place of the guide RNA. Assembly of these components at the *Per1* promoter allows the three effector domains (VP64, p65, and HSF1) to drive *Per1* transcription. To confirm the effectiveness of the CRISPR-SAM system, we transfected HT22 cells with all three plasmids, harvested the cells 24 or 48h later, and measured *Per1* mRNA expression. By 48h after transfection, *Per1* mRNA was significantly increased in the group given *Per1* sgRNA compared to controls **(Fig. 5E)**, confirming that the CRISPR-SAM system effectively drives *Per1* mRNA expression.

Next, 18-m.o. mice were given intra-hippocampal infusions of the CRISPR-SAM lentiviruses and trained in OLM 2 weeks later. As observed with our pLVX-v5Per1 overexpression, CRISPR-SAM-mediated *Per1* overexpression significantly improved memory performance in *Per1* sgRNA-infused mice relative to control animals (Fig. 5F) without affecting total exploration (Fig. 5G). Together, these results demonstrate that overexpression of *Per1* in the dorsal hippocampus is sufficient to ameliorate age-related impairments in long-term object location memory. *Per1* is therefore a key gene that is critical for long-term memory formation, is regulated by HDAC3, and is impaired in the aging brain.

## Discussion

Our results show that deletion or disruption of the repressive histone deacetylase HDAC3 ameliorates age-related impairments in both long-term memory and synaptic plasticity, consistent with the hypothesis that a repressive chromatin structure in aging neurons prevents normal gene expression required for long-term memory formation (Penner et al., 2010). Further, deletion of HDAC3 restores learning-induced expression of the circadian gene *Per1* in the dorsal hippocampus. As hippocampal PER1 expression is critical for long-term memory formation (Fig. 4) and overexpression of *Per1* in the hippocampus ameliorates age-related memory impairments (Fig. 5), PER1 is a potential mechanism through which deletion of HDAC3 improves memory and synaptic plasticity in aging mice. More broadly, age-related disruption of *Per1* might connect age-related impairments in both long-term memory and circadian rhythmicity, depending on the structure.

One key finding from the current study was that age-related impairments in hippocampal LTP could be ameliorated with HDAC3 deletion or disruption. This is consistent with recent work from the Sajikumar lab demonstrating that pharmacological blockade of HDAC3 can also ameliorate age-related impairments in associative hippocampal LTP (Sharma et al., 2015). Interestingly, we found that slices from old HDAC3^flox/flox^ and HDAC3(Y298H) animals failed to reach the same level of potentiation as slices from young HDAC3^flox/flox^ or HDAC3(Y298H) animals (Fig. 2C, 2F), suggesting that aging brains may have a lower plasticity ceiling than young brains. One possible explanation for this discrepancy is that age-related loss of synaptic contacts in CA1 (Barnes, 2003) could lower the plasticity ceiling in the aging hippocampus, as the fewer available synapses would become saturated more quickly than the relatively abundant synapses in the young DH. If this is the case, strengthening the stimulation protocol or providing spaced stimulation bouts (Kramar et al., 2012) should not further enhance LTP in the aging hippocampus, as no additional synapses are available. Further work will be necessary to determine the mechanism underlying this discrepancy.

Our RNA sequencing results demonstrated that only a small subset of genes fit the criteria of being “restored” in the aging brain by HDAC3 deletion (Fig. 3E). Thus, rather than recapitulating the gene expression profile of the young brain, deleting HDAC3 in the aging brain restores learning-induced expression of a few key genes that are critically important for long-term memory formation, including *Per1. Per1* appears to be directly regulated by HDAC3, as focal deletion of HDAC3 restored learning-induced *Per1* expression and HDAC3 is removed from the *Per1* promoter after learning. Further, *Per1* expression is necessary for learning, as siRNA-mediated knockdown of PER1 protein in the DH impaired long-term memory formation in young mice. This extends previous research showing that nonspecific deletion of *Per1* throughout the brain can impair memory formation in young mice (Jilg et al., 2010; Rawashdeh et al., 2014; Rawashdeh et al., 2016). Abnormal HDAC3-mediated repression of *Per1* in the aging brain is therefore a key event that could lead to age-related impairments in both long-term memory formation and circadian rhythmicity.

Circadian effects on long-term memory are traditionally believed to stem from dysregulation within the SCN, which then drives alterations in peripheral structures involved in memory formation, like the dorsal hippocampus. Little is known about the role of individual circadian clock genes in the DH, despite a clear connection between circadian rhythmicity and long-term memory formation. Memory is closely linked to time-of-day, as plasticity-related gene cascades show circadian oscillations (Eckel-Mahan et al., 2008; Rawashdeh et al., 2016) and memory can be acquired more easily at certain periods of the circadian cycle. For example, context fear conditioning is acquired more strongly during the daytime, when MAPK phosphorylation levels peak (Eckel-Mahan et al., 2008). Further, it is well-documented that aging is accompanied by a breakdown of circadian rhythms, presumably due to changes in the central circadian clock, the suprachiasmatic nucleus (SCN) (for review, Deibel et al., 2015). How this disruption in the circadian clock relates to the age-related impairments in memory is an open question. Our results suggest that HDAC3 limits learning-induced Per1 in the aging hippocampus, possibly contributing to the observed impairments in long-term memory. Epigenetic repression of Per1 may therefore represent an important interface between age-related impairments in both circadian rhythmicity and long-term memory formation.

Of the core canonical circadian clock genes, *Per1* is uniquely poised to dramatically affect hippocampal long-term memory. *Per1* appears to be predominantly involved in SCN output pathways and plays a key role in peripheral clocks downstream of the SCN, such as the hippocampus (Cermakian et al., 2001). Further, recent work suggests that *Per1* may “gate” spatial memory formation throughout the day/night cycle by controlling CREB phosphorylation (Jilg et al., 2010; Rawashdeh et al., 2014; Rawashdeh et al., 2016). Along with the results of the current study, this work indicates that PER1 expression is critically important for hippocampal long-term memory formation; reductions in PER1 that occur at night (Jilg et al., 2010) or with aging could impair hippocampal memory. To date, research implicating *Per1* in memory formation has exclusively relied on global knockouts that disrupt *Per1* expression in the core circadian clock and other regions in addition to memory-relevant structures like the hippocampus (Abarca et al., 2002; Jilg et al., 2010; Rawashdeh et al., 2014; Rawashdeh et al., 2016; Sakai et al., 2004), making it impossible to determine whether hippocampal PER1 is specifically required for memory formation. Indeed, global *Per1* deletion does affect circadian rhythmicity in some reports (Cermakian et al., 2001). Here, we show for the first time that reducing PER1 expression directly in the dorsal hippocampus can impair memory in young mice whereas local overexpression of *Per1* in the dorsal hippocampus can improve memory in aging mice.

Therefore, the core circadian clock gene *Per1* plays a key role within local memory structures to alter memory formation, even when SCN-based circadian rhythms are intact. More generally, this challenges the traditional hypothesis that circadian changes in memory formation are driven by alterations in the core circadian clock and instead supports the hypothesis that circadian clock genes play a more autonomous role in hippocampal cells, possibly “gating” memory formation based on the time of day (Rawashdeh et al., 2016).

## Author Contributions

J.L.K., D.P.M., and M.A.W. designed the experiments. J.L.K., Y.A., A.V.C., and D.P.M. conducted the experiments. J.L.K. and M.A.W. wrote the manuscript. E.K. conducted the electrophysiology and analyzed the results. J.L.K., E.M., and P.S.-C. designed and conducted the circadian rhythm experiment. Y.L., C.N.M., and P.B. performed the RNA sequencing analysis. A.J.L., A.O.W., G.S., D.R., and C.M.M. contributed to experiments. P.B., P.S.-C., and D.P.M. assisted in experimental design, data analysis, and manuscript preparation.

## Acknowledgements

We wish to thank all members of the Wood lab for scientific discussions and technical assistance. We would also like to acknowledge the University of California, Irvine Institute for Genomics and Bioinformatics High-Throughput Facility for help with RNA sequencing and Yuzo Kanomata for additional computing support. This work was supported by the National Institutes of Health grants (MH101491, AG051807, and AG050787 to M.A.W. and T32-AG000096-31 and F32-AG052303 to J.L.K.). The work of Y.L., C.M., and P.B. was in part supported by NSF grant IIS-1550705 and DARPA grant D17AP00002 to P.B. E.M. was supported by a long-term EMBO post-doctoral fellowship and work in the P.S.-C. lab was supported by grants from the National Institutes of Health, INSERM, and the Novo Nordisk Foundation Challenge Programme.

The authors declare no financial conflicts of interests. RNA sequencing data have been deposited in NCBI’s Gene Expression Omnibus and are accessible through GEO Series accession number GSE94832: https://www.ncbi.nlm.nih.gov/geo/query/acc.cgi?acc=GSE94832

## Methods

### Mice

Young adult mice were between 2-4 months old at the time of testing and aging mice were between 18-20 months old. Mice had free access to food and water and lights were maintained on a 12h light/dark cycle. All behavioral testing was performed during the light cycle. All experiments were conducted according to US National Institutes of Health guidelines for animal care and use and were approved by the Institutional Animal Care and Use Committee of the University of California, Irvine.

### Surgery

Mice were anesthetized with isoflurane (induced, 4%; maintained 1.5-2.0%) and placed in the stereotax. Injection needles were lowered to the dorsal hippocampus (AP, −2.0mm; ML, ±1.5mm, DV, −1.5mm relative to Bregma) as previously described (Kwapis et al., 2017a). 1.0μl of virus or siRNA was infused bilaterally into the DH at a rate of 6μl/hr. For the CRISPR-SAM lentiviral infusions, a cocktail of the three viruses was infused to a final volume of 1.5μL per hemisphere at the same rate. Viral infusions were performed 2 weeks before behavioral analysis whereas siRNA knockdown was performed 2d before training(McQuown et al., 2011). For all injection experiments animals were randomly assigned to the different injection conditions (with the exception of HDAC3^+/+^ and HDAC3^flox/flox^ mice, which were all injected with AAV-CaMKII-Cre). For all behavioral experiments, animals within each viral condition were randomly assigned to homecage/trained groups and all conditions (objects, boxes, etc.) were counterbalanced between groups.

### AAV Production

AAV2.1-CaMKII-Cre was purchased from Penn Vector Core (titer: 1.81 × 10^13^ GC/mL). For AAV2.1-HDAC3(Y298H)-v5, we amplified wildtype HDAC3 from hippocampal cDNA and cloned the product into a modified pAAV-IRES-hrGFP (Agilent) under control of the CMV promoter and β-globin intron. The 3x-FLAG tag, IRES element, and hrGFP were removed from the vector and replaced with a V5 tag, allowing for a C-terminal fusion to HDAC3 (plasmid MW91). To create the point mutation, we changed the nucleotides to code for a histidine residue in place of tyrosine at amino acid 298 (plasmid MW92). For the empty vector control, the HDAC3 coding sequence was not present, but all other elements remain (plasmid MW87). AAV was made by the Penn Vector Core and the final titers were determined by qPCR (AAV-HDAC3(Y298H): 6.48 × 10^12^ GC/mL; AAV-EV: 1.35 × 10^13^ GC/mL).

### Lentivirus Production

For the CRISPR/dCas9 Synergistic Activation Mediator (SAM) system, lentiviral plasmids were purchased from Addgene for the dCas9-VP64 (#61422-LVC) and MS2-P65-HSF1 (#61426-LVC) constructs (titers ≥ 8×10^6^ TU/mL). For the *Per1* sgRNA, we cloned and inserted an antisense guide sequence corresponding to the CRE element in the *Per1* promoter (AGAGGGAGGTGACGTCAAAG) into the Addgene sgRNA(MS2) cloning backbone (#61427). The empty vector control was identical, except that no guide sequence was cloned into the plasmid. Lentiviruses for both the *Per1* sgRNA (titer: 6.8 × 10^7^ IFU/mL) and EV sgRNA (3.5 × 10^7^ IFU/mL) were produced by the USC School of Pharmacy Lentiviral Laboratory.

For the pLVX-V5Per1 overexpression construct, we amplified full-length wildtype *Per1* from the Addgene pCMV-Sport2-mPer1 plasmid (#16203) and cloned the product into a modified pLVX-EF1α-IRES-mCherry backbone (Takara, #631987). The IRES and mCherry elements were removed and were replaced with a V5 tag, allowing for an N-terminal fusion to PER1 (plasmid MW206). For the empty vector (EV) control, the PER1 coding sequence was not present but all other elements remained (Plasmid MW93). Lentiviruses for the pLVX-V5Per1 (titer: 1.3 × 10^8^ IFU/mL) and pLVX-EV (titer: 1.5 × 10^8^ IFU/mL) were produced by the USC School of Pharamacy Lentiviral Laboratory. All lentiviral constructs were expressed under the EF1α promoter.

### Cell Culture Verification of CRISPR/dCas9 SAM System

To verify that the CRISPR-SAM system can effectively drive *Per1* transcription, HT22 cells were transfected with dCas9-VP64 (lenti MS2-P65_HSF1_Blast was a gift from Feng Zhang Addgene plasmid #61425), MS2-P65-HSF1 (lenti dCAS-VP64_Blast was a gift from Feng Zhang, Addgene plasmid #61426), and either *Per1* sgRNA(MS2) (Per1) or the non-targeting empty vector sgRNA(MS2) (EV) (lenti sgRNA (MS2)_zeo backbone was a gift from Feng Zhang, Addgene plasmid #61427) using Lipofectamine LTX (Invitrogen). Cells were harvested after 24h or 48h, lysed, and mRNA was isolated as described above. qRT-PCR was performed as described above using the *Per1* primers and probe listed in Table S4.

### siRNA

For the *Per1* knockdown experiment, Accell SMARTpool small interfering RNAs (siRNAs; Dharmacon, GE) targeting *Per1* were diluted to a final total concentration of 10 μM in ddH20 and infused into the DH (1.0 μ1/side). Accell non-targeting pool siRNA was used as the control (total concentration, 10 μM) and was infused in the same manner. For siRNA experiments, mice were handled and habituated as described above and surgery was performed the day after the final day of habituation. Mice were given a full day of recovery after surgery and were trained the following day (~48h after surgery) to ensure maximal target knockdown. To ensure knockdown, mice were sacrificed ~1h after test and punches from the dorsal hippocampus were processed with western blots to ensure knockdown of PER1 protein.

### Object Location and Object Recognition Memory Tasks

Object location and object recognition memory tasks were performed as previously described(Vogel-Ciernia et al., 2013; Vogel-Ciernia and Wood, 2014). Mice were handled for 2 minutes/day for 4d and then habituated to the context for 5 minutes/day for 6 consecutive days in the absence of objects. During training, mice were exposed to two identical objects (100mL beakers, spice tins, or glass candle holders) and allowed to explore for 10 minutes. During the retention test (24h later for long-term memory or 60m later for short-term memory), mice were allowed to explore for 5 minutes. For object location memory, one of the two familiar objects was moved to a new location. For object recognition memory, the object locations remained constant but one of the objects was replaced with a new item. Habituation for object recognition memory began at least one week after the completion of OLM testing and a new context and unfamiliar objects were used(Vogel-Ciernia and Wood, 2014). Exploration was scored when the mouse head oriented toward the object and came within 1cm or when the nose touched the object. Total exploration time was recorded (*t*) and preference for the novel item was expressed as a discrimination index (*DI*=(*t*_novel_ = *t*_familiar_) × 100%). Mice that explored both objects for less than 2s during testing or 3s during training were removed from further analysis. Mice that showed a preference for one object during training (DI > ±20) were also removed. Habituation sessions were analyzed (to determine distance traveled and speed) using ANY-maze behavioral analysis software (Stoelting Co). All habituation, training, and testing were performed by experimenters blinded to the experimental groups.

### Context fear conditioning

Contextual fear conditioning was performed as previously described(Kwapis et al., 2017a; Vogel-Ciernia et al., 2013). Mice were handled for 5d before conditioning. During acquisition, mice were exposed to the context for 2 minutes and 28s followed by a 2s (0.75mA) shock. Mice remained in the contxt for an additional 30s before being removed. Mice were sacrificed 60m after training along with homecage controls that were handled but not trained. Freezing behavior was measured using Ethovision 11 software (Noldus)(Kwapis et al., 2017a).

### Elevated plus maze

The plus-maze was conducted as previously described(Vogel-Ciernia et al., 2013) by an experimenter blind to the experimental groups. One week after the completion of ORM, a subset of mice were tested on the plus maze. Two arms of the maze were open (30 × 5 cm) and two arms were enclosed (30 × 5 × 15 cm), connected by a central platform (5 × 5 cm). The maze was elevated 40cm above the floor. Mice were tested for 5 min on the apparatus, which consisted of placing each mouse onto the central platform facing one of the open arms. Between subjects, the maze was cleaned with 70% ethanol. The percentage of time spent in the closed and open arms was scored using ANY-maze software.

### Circadian rhythm analysis

Young (3-m.o) and aging (18-m.o.) HDAC3^+/+^ and HDAC3^flox/flox^ mice were bred and housed under a 12h light/dark (LD) cycle. 2 weeks after AAV-CaMKII-Cre infusion (described above), mice were transferred to an isolated 12h LD entrainment room for 7d. Mice were then transferred into a light-protected activity analysis room, where locomoter activity was analyzed using optical beam motion dectors (Philips Respironics). Activity was monitored during 2 weeks of LD cycle entrainment in the light-protected room before mice were switched to constant darkness (DD) for an additional 3 weeks. Activity monitoring continued throughout the DD phase to determine whether HDAC3 deletion in the DH affected endogenous circadian rhythms. Data were collected using Minimitter VitalView 5.0 and Clocklab software (Actimetrics) was used to determine the onset of free activity. Tau values were calculated by obtaining the slope of this onset and calculating the least-squares fit with Clocklab software(Eckel-Mahan and Sassone-Corsi, 2015; Orozco-Solis et al., 2016).

### Immunohistochemistry

Immunofluorescence was conducted as previously described (Kwapis et al., 2017a; Lopez et al., 2016; White et al., 2016). After the completion of behavior, mice were euthanized by cervical dislocation and their brains were removed and flash-frozen in ice-cold isopentane. 20μm slices were collected throughout the dorsal hippcampus, thaw-mounted on slides, and stored at −80°C. Slides were fixed with with 4% paraformaldehyde (10-min), permeabilized in 0.01% Triton X-100 in 0.1 M PBS (5-min), and blocked for 1h with 8% normal goat serum (Jackson). Slides were incubated overnight (4°C) in rabbit antibody to HDAC3 (1:250, Abcam, ab32369) or V5 (1:1000, Abcam, ab9116). The following day, slides were washed and incubated for 1h at room tempoerature with goat anti-rabbit Alexa 488 (1:1000, Invitrogen) in the dark. Slides were then washed with PBST and incubated for 50m in NeuroTrace 530/615 (1:50; ThermoFisher), a fluorescent nissl stain. To quench nonspecific autofluorescence(Schnell et al., 1999), slices were then washed in PBS with 0.01% Triton, rinsed in water, and incubated for 10-min in 10mM CuSO_4_ in 50mM ammonium acetate buffer. Slices were again rinsed in water, washed in PBS and coverslipped with VectaShield Antifade mounting medium (Vector Laboratories).

All images were acquired with an Olympus Scanner VS110 with a 20× apochromatic objective (numerical aperture 0.75) with VS110 scanner software. All treatment groups were represented on each slide and all images on a slide were captured with the same exposure time under nonsaturating conditions. Immunolabeling intensity was quantified with ImageJ by sampling the optical density of the cell layer in CA1 and subtracting a sample of background fluorescence in the same image. For all AAV experiments, animals that failed to express the virus in area CA1 of the dorsal hippocampus were excluded from analyses. Imaging and quantification was performed by experimenters blind to the experimental conditions.

### Western blot

To verify PER1 knockdown, Per1 siRNA and control siRNA mice were sacrificed 1h after testing and brains were flash-frozen. Brains were coronally sectioned and 1mm DH punches were collected from 500 μm slices. Punches were homogenized in T-PER buffer (Thermo Fisher) with Halt protease and phosphatase inhibitor (Thermo Fisher) using a dounce homogenizer. Protein lysates were quantified using a modified Bradford assay (BioRad) and 10 μg (for HDAC3 and phospho-HDAC3) or 50 μg (for PER1) total protein lysate was loaded into each lane of a 7.5% NuPAGE Bis-Tris gel (Thermo Fisher). Gels ran for 50 minutes at 200 volts and blots were transferred overnight at 15V at 4°C onto nitrocellulose membranes (Novexi, LC2001). The following day, membranes were incubated in blocking buffer for 1h (5% nonfat milk in Tris-buffered saline with 0.01% Tween 20), washed (0.1% Tween 20 in TBS) and then incubated in primary antibody (1:500, rabbit anti-Per1, Thermo Fisher) in primary antibody buffer (3% BSA in TBS with 0.1% Tween) overnight at 4°C. The membranes were then washed and incubated in HRP-conjugated mouse antibody to rabbit (1:10,000, Jackson Laboratories) for 1h. Membranes were washed and developed using Pierce SuperSignal West Pico Chemilumenescent Substrate (Pierce, 34077). Multiple film exposures were used to verify linearity. Blots were washed and stripped for 10m with Restore Western Blot Stripping Buffer (Thermo Fisher #21059), washed again, and then re-probed overnight (4°C) with a rabbit antibody to GAPDH (1:1000, Santa Cruz Biotechnology, SC25778). Relative optical densities were calculated from scanned film using ImageJ (US National Institutes of Health) by an experimenter blind to the experimental conditions. All values were normalized to GAPDH expression levels.

### qRT-PCR

Quantitative real-time RT-PCR was performed as previously described (Kwapis et al., 2017a; Vogel-Ciernia et al., 2013). Tissue was collected from DH punches (described above) and frozen at −80°C until processing. RNA was isolated using an RNeasy Minikit (Qiagen, #74104) and cDNA was created using the Transcriptor First Strand cDNA Synthesis kit (Roche, 04379012001). Primers and probes were derived from the Roche Universal ProbeLibrary (Table S4) and were used for multiplexing in the Roche Light-Cycle 480 II machine (Roche). All values were normalized to *Gapdh.* Analyses and statistics were performed using the Roche proprietary algorhithms and REST 2009 software based on the Pfaffl method (Pfaffl, 2001; Pfaffl et al., 2002).

### RNA sequencing

RNA was isolated from dorsal hippocampus punches as described above, using the RNeasy minikit (Qiagen, 74104). RNA quality was assessed by Bioanalyzer and samples with an RNA integrity number >9 were included in our analysis. cDNA libraries for each group were prepared as described previously(Vogel-Ciernia et al., 2013), according to the TruSeq RNA Sample Preparation Guide (Illumina). Briefly, 250ng of total RNA from each mouse was purified with poly-T oligo-attached magnetic beads and heat fragmented. The first and second strand cDNA were then synthesized and purified. After the ends were blunted, the 3’ end was adenylated to prevent concatenation of the template during adaptor ligation. For each group, a unique adaptor set was added to the ends of the cDNA and the libraries were amplified by PCR. The quality of the library was assessed by Bioanalyzer and quantified using qRT-PCR with a standard curve prepared from a commercial sequencing library (Illumina). Samples were mutliplexed, with each behavioral group represented in each flow cell of the sequencer. 10nM of each library was pooled in four multiplex libraries and sequenced on an Illumina HiSeq 2500 instrument during a single-read 50 bp sequencing, run by the Genomice High-Throughput Facility at the University of California, Irvine. The resulting sequencing data for each library were post-processed to produce FastQ files. The data were then demultiplexed and filtered using Illumina CASAVA 1.8.2 software as well as in-house software. Poor-quality reads (failing Illumina’s standard quality tests) and control reads successfully aligned to the PhiX control genome were removed from analyses. The quality of the remaining sequences was further assessed using PHRED quality scores produced in real time during the base-calling step of the sequencing run (Supplementary Figure 3a).

### Alignment to the reference genome and transcriptome

The reads from each replicate experiment were separately aligned to the reference genome and corresponding transcriptome using short-read aligners ELAND v2e (Illumina) and Bowtie(Langmead et al., 2009). Reads uniquely aligned by both tools to known exons or splice junctions with no more than two mismatches were included in the transcriptome. Reads uniquely aligned, but with more than two mismatches were removed from analyses. Similarly, reads matching several locations in the reference genome were removed from analysis. The percentage of reads assigned to the reference genome and transcriptome using this protocol is reported for each group of replicates (Supplementary Figure 3b).

### Gene expression and differential analysis

Gene expression levels were directly computed from the read alignment results for each replicate. Standard RPKM values(Mortazavi et al., 2008), reads per kilobase of exon model per million mapped reads) were extracted for each gene covered by the sequencing data and each replicate used in this study.

Differential transcriptional analyses were performed using Cyber T(Baldi and Long, 2001; Kayala and Baldi, 2012) across each pair of groups (homecage versus 60m after training) to identify genes up- or down-regulated after learning. In addition to the 18-month-old HDAC3^+/+^ and HDAC3^flox/flox^ mice trained for the current study, identically processed RNA-sequencing data from 3-month old C57 wildtype mice (homecage and OLM-trained in the same manner as the current study) from a previous study(Vogel-Ciernia et al., 2013) was used for differential analyses. The p-value threshold used for determining differential expression is 0.05 for all groups. The sets of genes upregulated after learning (compared to homecage) for these 3 groups (young wildtype, aging HDAC^+/+^, or aging HDAC3^flox/flox^) were intersected to determine genes common to two or more groups. Enrichment of each group for tissue-specific expression, Gene Ontology terms(Ashburner et al., 2000), and KEGG pathways(Kanehisa et al., 2012; Ogata et al., 1999) was assessed using DAVID(Dennis et al., 2003), based on differentially expressed genes after learning. Data visualization was performed using ‘matplotlib’ for python and ‘ggplot’ for R.

### Chromatin immunoprecipitation

ChIP was performed on DH punches as previously described(Kwapis et al., 2017a; Malvaez et al., 2013; Rogge et al., 2013), based on the protocol from the Millipore ChIP kit. Tissue was cross-linked with 1% formaldehyde (Sigma), lysed and sonicated, and chromatin was immunprecipitated overnight with 2 μl of anti-H4K8Ac (Millipore #17-10099) or 4 μl of anti-HDAC3 (Millipore #17-10238) or an equivalent amount of Normal Rabbit Serum (H4K8Ac negative control, Millipore) or anti-mouse IgG (HDAC3 negative control, Millipore). After washing, chromatin was eluted from the beads and reverse cross-linked in the presence of proteinase K before column purification of DNA. Primer sequences for the promoters of each gene were designed by the Primer 3 program (Table S4). Five μl of input, anti-H4K8Ac IgG/anti-HDAC3 IgG, or anti-rabbit/mouse IgG immunoprecipitate from each animal were examined in duplicate. To normalize ChIP-qPCR data, we used the percent input method. The input sample was adjusted to 100% and both the IP and IgG samples were calculated as a percent of this input using the formula 100*AE^(adjusted input – Ct(IP)). Fold enrichment was then calculated as a ratio of the ChIP to the average IgG. An in-plate standard curve determined amplification efficiency (AE).

### Slice preparation and recording

Young (approximately 3-m.o.) and aging (18-m.o.) mice were stereotaxically infused with virus. For the first experiment, young and old HDAC3^+/+^ and HDAC3^flox/flox^ mice were infused with AAV-CaMKII-Cre bilaterally into the DH. For the second experiment, young and old wildtype C57 mice were infused with AAV-HDAC3(Y298H) into the DH of one hemisphere and AAV-EV control into the other hemisphere. Two weeks after infusion (to allow for optimal virus expression)^2^, transverse hippocampal slices (320 μm) were prepared as previously described (Haettig et al., 2011; Vogel-Ciernia et al., 2013) and placed in an interface recording chamber with preheated (31 ± 1°C) artificial cerebrospinal fluid (124 mM NaCl, 3mM KCl, 1.25mM KH_2_PO_4_, 1.5 mM MgSO_4_, 2.5 mM CaCl_2_, 26 mM NaHCO_3_, and 10 mM _D_-glucose). Slices were continuously perfused at a rate of 1.75-2 ml/min while the surface of the slices was exposed to warm, humidified 95% O_2_ / 5% CO_2_. Recordings began after at least 2h of incubation.

Field excitatory postsynaptic potentials (fEPSPs) were recorded from CA1b stratum radiatum using a single glass pipette (2-3MΩ) filled with 2 M NaCl. Stimulation pulses (0.05 Hz) were delivered to Schaffer collateral-commissural projections using a bipolar stimulating electrode (twisted nichrome wire, 65 μm) positioned in CA1c. Current intensity was adjusted to obtain 50% of maximal fEPSP response. After a stable baseline was established, LTP was induced with a single train of five “theta” bursts, in which each burst (four pulses at 100Hz) was delivered 200 ms apart (i.e. at theta frequency). The stimulation intensity was not increased during TBS. Data were collected and digitized by NAC 2.0 Neurodata Acquisition System (Theta Burst).

### Statistical analysis

Statistical analyses were conducted as indicated in the text and figure legends using Prism 6 (GraphPad). Our analytic approaches are based on previously published work(Kwapis et al., 2017a; Lopez et al., 2016; Vogel-Ciernia et al., 2013; White et al., 2016). No statistical methods were used to predetermine sample sizes, but our sample sizes are similar to those generally used in the field, including those reported in previous publications(Haettig et al., 2011; Kwapis et al., 2017a; Lopez et al., 2016; Vogel-Ciernia et al., 2013). Data distribution was assumed to be normal, with similar variance expected among groups, but this was not formally tested. When an experiment had two groups to compare, Student’s t-tests were used. When two factors where compared, (such as age and genotype), data were analyzed with two-way ANOVAs followed by Sidak’s multiple comparisons *post hoc* tests. All analyses are two-tailed, with an α value of 0.05 required for significance. Error bars in all figures represent S.E.M. For all experiments, values ±2SD from the group mean were considered outliers and were removed from analyses.

### Data availability

The data supporting the findings of this study are available from the corresponding author upon reasonable request.

**Supplementary Figure 1.**
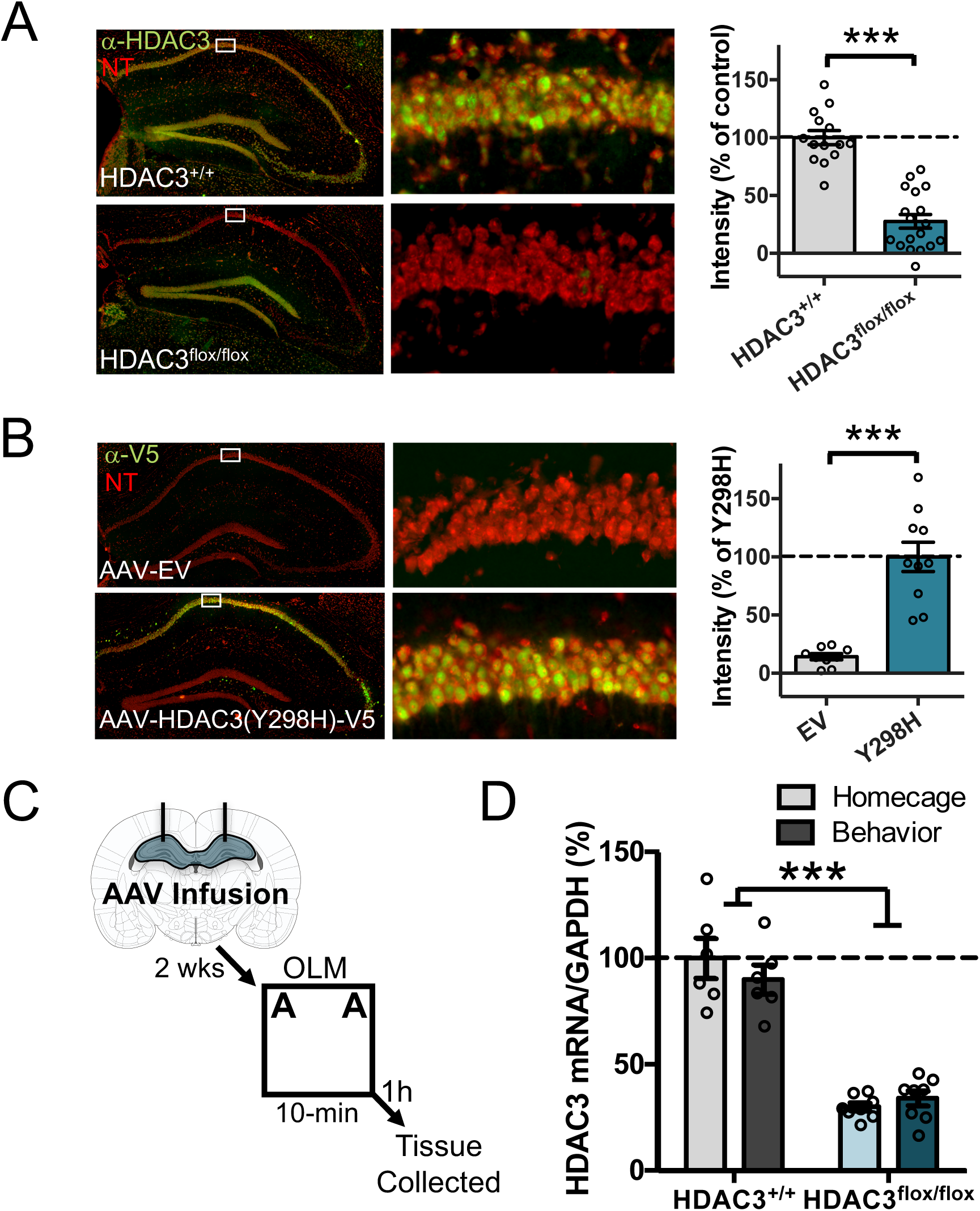
Genetic deletion or activity-specific disruption of HDAC3 in the dorsal hippocampus of aging mice. **(A)** Representative immunofluorescence images of HDAC3 (green) expression in HDAC3^+/+^ and HDAC3^flox/flox^mice injected with AAV2.1-CaMKII-Cre. Cell bodies were counterstained with a fluorescent Nissl stain (NeuroTrace, red). **Right:** mean intensity of HDAC3 immunofluorescence sampled from CA1 (normalized to background). HDAC3 expression was significantly reduced in HDAC3^flox/flox^mice compared to HDAC3^+/+^controls (*t*(30)=8.45, \*\*\**p*<0.0001, *n*=14(5F), 18(6F)). **(B)** Representative immunfluorescence images of V5 (green) expression in wildtype mice injected with AAV-EV (top) or AAV2.1-CMV-HDAC3(Y298H)-V5 (bottom) counterstained with NeuroTrace (red). **Right:** Mean intensity of V5 immunofluorescence (normalized to background). V5 expression was significantly higher in mice injected with AAV-HDAC3(Y298H)-V5 compared to AAV-EV controls (*t*(17)=6.33, *p*<0.0001, *n*=9,10, all male). **(C)** Experimental procedure. **(D)** HDAC3 mRNA expression in the dorsal hippocampus. HDAC3^+/+^ (gray bars) or HDAC3^flox/flox^(blue bars) mice infused with AAV-Cre were sacrificed at rest (homecage) or 60m after OLM (OLM). HDAC3 mRNA expression was significantly reduced in HDAC3^flox/flox^ animals compared to HDAC3^+/+^ animals regardless of whether they received OLM training (Genotype group difference only, *F*(1,24) = 132.1,\*\*\**p*<0.0001, n=6(2F), 6(3F), 8(5F), 8(5F)).

**Supplementary Figure 2.**
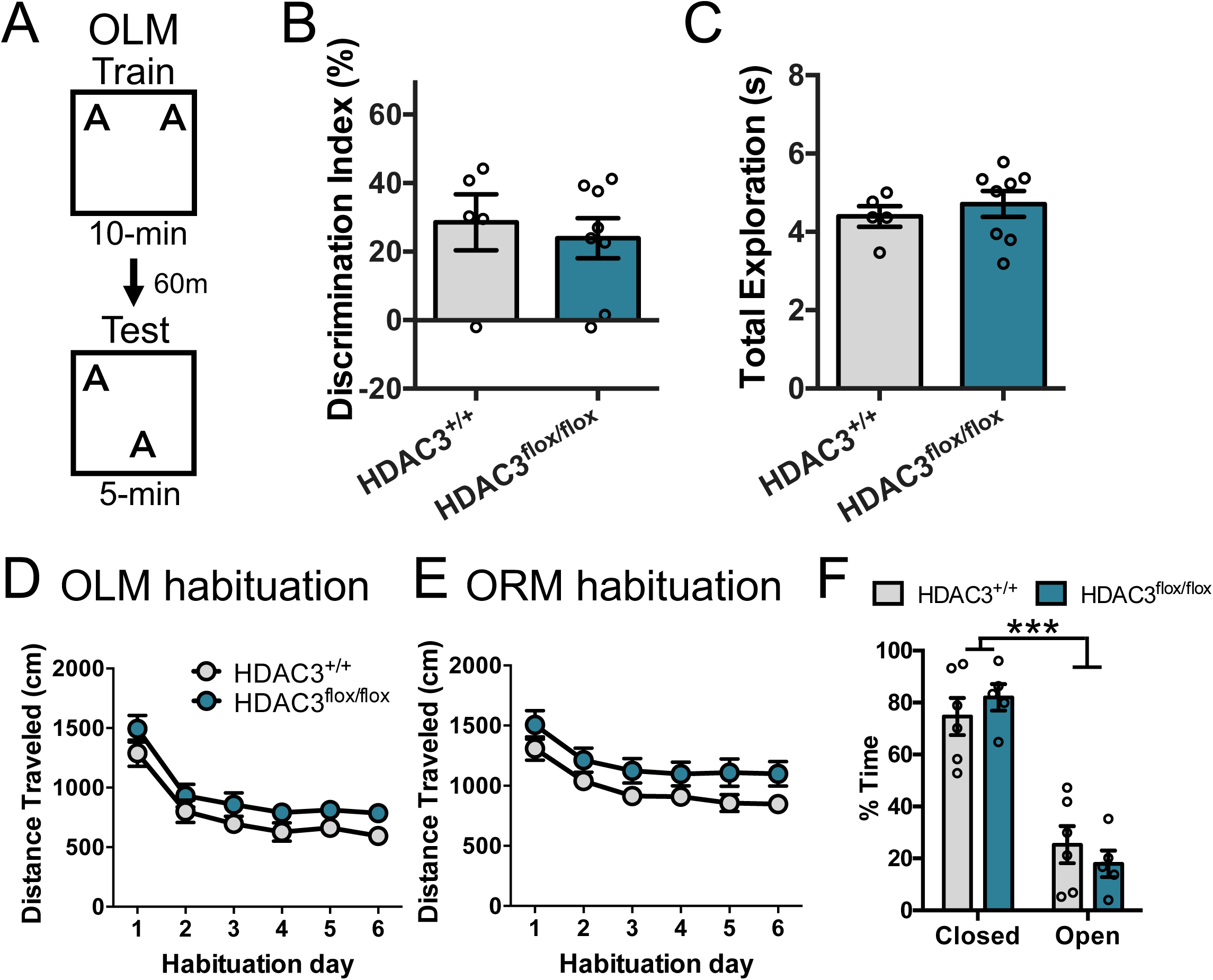
All groups show normal short-term memory, movement, and anxiety. **(A)** Experimental schematic testing short-term memory. Two weeks after AAV-CaMKII-Cre infusion into the DH, 18-month-old HDAC3^+/+^ and HDAC3^flox/flox^mice were trained in OLM and tested 60m later. **(B)** Both groups showed intact memory for OLM when tested 60m after training. No significant differences were observed between groups (*t*_(11)_=0.477, *p*=0.643; n=5,8; all male). **(C)** Both groups also showed similar levels of total exploration (*t*_(11)_=0.68, *p*=0.511). **(D)** Total distance traveled during each day of OLM habituation for mice in Figure 1B. No group differences were observed and individual comparisons between groups each day revealed no significant differences. **(E)** Total distance traveled during ORM habituation for mice in Figure 1G. No group differences were observed and individual comparisons between groups each day revealed no significant differences. **(F)** A subset of animals from Figure 1B were tested in the elevated plus maze. Both groups showed similar levels of anxiety, as indicated by the percentage of time spent in the closed and open arms (Effect of Arm only, *F*_(1,9)_=38.47, \*\*\**p*<0.001, n= 6(4R), 5(1 F)).

**Supplementary Figure 3.**
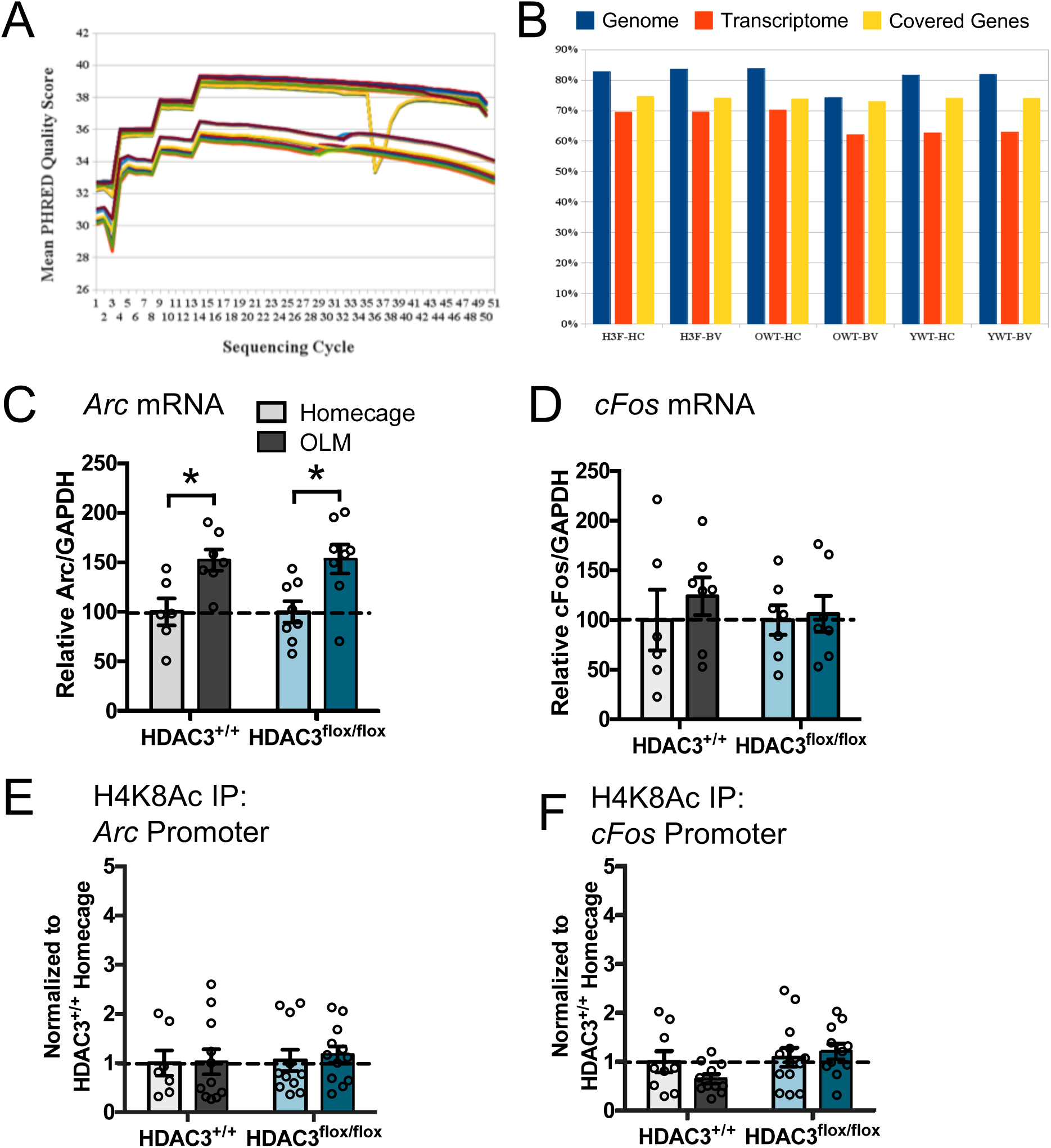
RNA sequencing quality controls and characterization of individual genes. **(A)** Mean PHRED quality scores for each animal in the six different groups: 18-m.o. HDAC3^flox/flox^ homecage (H3F-HC), 18-m.o. HDAC3^flox/flox^ behavior (H3F-BV), 18-m.o. wildtype homecage (OWT-HC), 18-m.o. wildtype behavior (OWT-BV), 3-m.o. wildtype homecage (YWT-HC), and 3-m.o. wildtype behavior (YWT-BV). **(B)** Percentage of short-reads uniquely aligned to the reference genome (blue bars) and transcriptome (red bars) and percentage of annotated genes covered by the sequencing reads (yellow bars) for each of the six groups. **(C-F)** Characterization of individual genes in 18-m.o. wildtype and HDAC3^flox/flox^mice after learning, from Figure 3. **(C)** *Arc* mRNA expression (Effect of training only *F*_(1,25)_=17.31, p = 0.0003, n=6(2F),7 (3F),8(5F), 8(5F)). **(D)** *cFos* mRNA expression (No significant effects, n=6(2F), 7(3F), 7(4F), 7(5F)). **(E)** H4K8Ac occupancy at the *Arc* promoter (No significant effects, n = 7(5F), 11 (4F), 11 (4F), 12(7F)). **(F)** H4K8Ac occupancy at the *cFos* promoter (No significant effects, n=9(5F), 11 (4F), 13(6F), 11 (6F)).

**Supplementary Figure 4.**
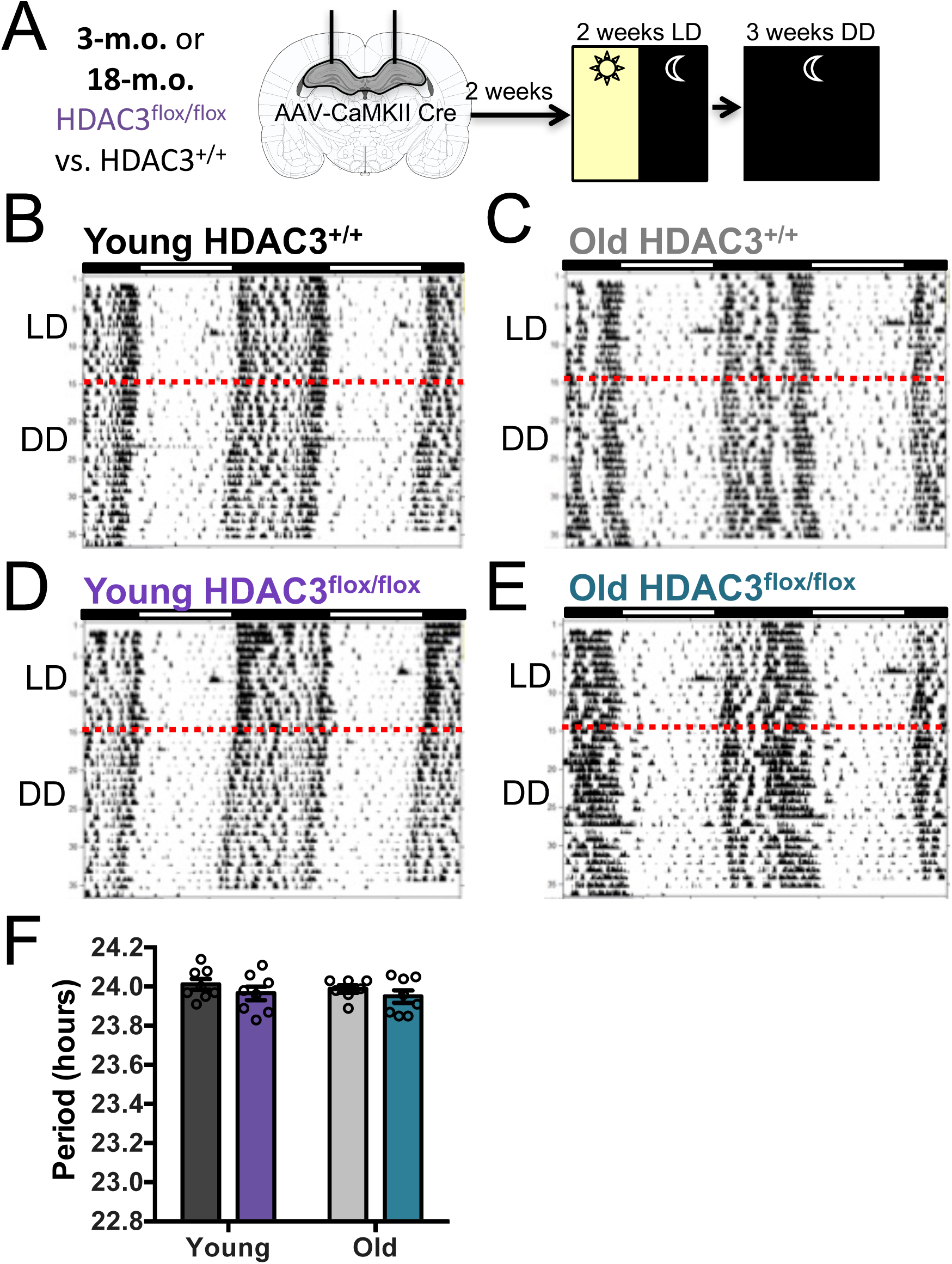
Deletion of HDAC3 in the DH does not affect circadian rhythm of young or old mice. **(A)** Experimental procedure. **(B-E)** Representative actograms depicting circadian behavior for each group. Red dotted line indicates the switch from LD to DD. **(B)** Young HDAC3^+/+^ actogram. **(C)** Old HDAC3^+/+^ actogram. **(D)** Young HDAC3^flox/flox^ actogram. **(E)** Old HDAC3^flox/flox^ actogram. **(F)** HDAC3 deletion in DH did not affect free-running tau in either young or old mice (no significant effects, n=8,8,7,8; all males).

**Table S1.** Genes with increased (top) or decreased (bottom) expression in young (3-m.o.) wildtype mice 60m after OLM training (trained) compared to baseline (homecage). The top 20 genes increased or decreased (based on *p*-value, homecage vs trained) are shown.

| Increased expression |  |  |  |  |  |
| --- | --- | --- | --- | --- | --- |
|  | Gene | Homecage<br>(Mean) | Trained<br>(Mean) | Fold Change<br>(log) | p-value |
| 1 | AF357426 | 0.123 | 1.062 | 0.939 | 9.75919E-12 |
| 2 | A230065H16Rik | 0.651 | 3.996 | 3.345 | 1.28698E-10 |
| 3 | Glp1r | 0.227 | 1.091 | 0.864 | 2.66057E-08 |
| 4 | Prdm12 | 0.072 | 0.456 | 0.383 | 2.43083E-07 |
| 5 | Gh | 0.007 | 0.350 | 0.343 | 2.02023E-06 |
| 6 | Ano2 | 0.389 | 1.077 | 0.688 | 3.13868E-06 |
| 7 | Sln | 0.668 | 2.403 | 1.734 | 3.40852E-06 |
| 8 | Igfbpl1 | 1.597 | 2.459 | 0.862 | 5.53357E-06 |
| 9 | Ctxn3 | 0.943 | 2.588 | 1.646 | 6.99663E-06 |
| 10 | Ngb | 0.552 | 1.313 | 0.761 | 8.55593E-06 |
| 11 | Magel2 | 0.141 | 0.418 | 0.277 | 2.82686E-05 |
| 12 | Mir702 | 0.000 | 0.110 | 0.110 | 3.61675E-05 |
| 13 | Isl1 | 0.018 | 0.239 | 0.221 | 5.03269E-05 |
| 14 | C130060K24Rik | 0.051 | 0.230 | 0.179 | 7.6283E-05 |
| 15 | Gpr88 | 0.762 | 2.072 | 1.310 | 0.000132281 |
| 16 | Mir7009 | 0.000 | 0.158 | 0.158 | 0.000179688 |
| 17 | Zic1 | 2.590 | 7.161 | 4.571 | 0.000236958 |
| 18 | Sost | 0.056 | 0.253 | 0.197 | 0.000278343 |
| 19 | C130074G19Rik | 5.293 | 8.409 | 3.116 | 0.000297396 |
| 20 | Dlx1 | 4.424 | 6.604 | 2.180 | 0.000537434 |
| Decreased expression |  |  |  |  |  |
|  | Gene | Homecage<br>(Mean) | Trained<br>(Mean) | Fold Change<br>(log) | p-value |
| 1 | Mir804 | 0.380 | 0.065 | -0.316 | 4.79591E-08 |
| 2 | DQ267100 | 1.702 | 0.561 | -1.141 | 5.03473E-06 |
| 3 | Mir486 | 0.312 | 0.058 | -0.254 | 7.96871E-06 |
| 4 | Cpne7 | 47.046 | 38.314 | -8.732 | 8.44389E-06 |
| 5 | Mir1668 | 0.447 | 0.064 | -0.383 | 1.00325E-05 |
| 6 | 2310003N18Rik | 0.039 | 0.000 | -0.039 | 2.98682E-05 |
| 7 | D16Ertd519e | 0.029 | 0.000 | -0.029 | 0.000129378 |
| 8 | Mir698 | 0.066 | 0.000 | -0.066 | 0.000159265 |
| 9 | 1700020M21Rik | 0.028 | 0.000 | -0.028 | 0.000168869 |
| 10 | Mir367 | 0.039 | 0.000 | -0.039 | 0.000393218 |
| 11 | Grp | 8.839 | 6.777 | -2.061 | 0.000451035 |
| 12 | Eif3j1 | 0.445 | 0.000 | -0.445 | 0.001197766 |
| 13 | Snord123 | 0.136 | 0.000 | -0.136 | 0.001221741 |
| 14 | Hbq1a | 0.295 | 0.068 | -0.227 | 0.001656455 |
| 15 | Mir202 | 0.029 | 0.000 | -0.029 | 0.002539309 |
| 16 | Nov | 37.795 | 30.697 | -7.097 | 0.002754578 |
| 17 | Snora34 | 1.884 | 0.725 | -1.160 | 0.002813874 |
| 18 | Mir669n | 0.090 | 0.000 | -0.090 | 0.003749518 |
| 19 | Mir6947 | 0.080 | 0.014 | -0.065 | 0.004189413 |
| 20 | Camp | 0.057 | 0.000 | -0.057 | 0.004231699 |

**Table S2.** Genes with increased (top) or decreased (bottom) expression in old (18-m.o.) wildtype mice 60m after OLM training (trained) compared to baseline (homecage). The top 20 genes increased or decreased (based on *p*-value, homecage vs trained) are shown.

| Increased expression |  |  |  |  |  |
| --- | --- | --- | --- | --- | --- |
|  | Gene | Homecage<br>(Mean) | Trained<br>(Mean) | Fold Change<br>(log) | p-value |
| 1 | Nr4a3 | 13.257 | 20.963 | 7.706 | 7.43057E-06 |
| 2 | Ak4 | 6.212 | 9.271 | 3.059 | 8.98888E-05 |
| 3 | Mir344d-2 | 0.000 | 0.249 | 0.249 | 0.000105067 |
| 4 | Slit2 | 1.970 | 3.727 | 1.757 | 0.000166913 |
| 5 | Tmem114 | 2.091 | 3.443 | 1.351 | 0.000181875 |
| 6 | Mir7054 | 0.009 | 0.261 | 0.252 | 0.000278494 |
| 7 | Sgk1 | 24.420 | 37.819 | 13.399 | 0.000317729 |
| 8 | Mir7665 | 0.000 | 0.098 | 0.098 | 0.000451994 |
| 9 | Mir7232 | 0.009 | 0.289 | 0.280 | 0.000463299 |
| 10 | Gm11213 | 0.000 | 0.105 | 0.105 | 0.000771559 |
| 11 | Sap18 | 1.153 | 2.875 | 1.722 | 0.000831838 |
| 12 | Lpl | 10.900 | 15.275 | 4.374 | 0.001252228 |
| 13 | Cldn22 | 2.520 | 4.334 | 1.814 | 0.001502037 |
| 14 | Cox8b | 0.073 | 7.384 | 7.311 | 0.001810557 |
| 15 | Pvrl3 | 6.285 | 9.501 | 3.216 | 0.001872061 |
| 16 | Ccdc3 | 7.139 | 10.598 | 3.459 | 0.001908397 |
| 17 | Coch | 1.422 | 4.054 | 2.632 | 0.001920303 |
| 18 | Arl4d | 7.804 | 11.689 | 3.885 | 0.002276406 |
| 19 | Mir7662 | 0.136 | 0.682 | 0.546 | 0.002427375 |
| 20 | Pon1 | 0.000 | 0.509 | 0.509 | 0.002437193 |
| Decreased expression |  |  |  |  |  |
|  | Gene | Homecage<br>(Mean) | Trained<br>(Mean) | Fold Change<br>(log) | p-value |
| 1 | Cd72 | 31.100 | 4.614 | -26.487 | 9.05719E-07 |
| 2 | Irf8 | 24.878 | 8.112 | -16.766 | 1.64456E-06 |
| 3 | Il4ra | 10.307 | 2.185 | -8.122 | 1.74389E-06 |
| 4 | Fcgr4 | 28.396 | 5.438 | -22.958 | 2.21299E-06 |
| 5 | Themis2 | 9.935 | 2.454 | -7.481 | 2.38824E-06 |
| 6 | Snord43 | 0.445 | 0.000 | -0.445 | 2.52343E-06 |
| 7 | Klrb1b | 1.652 | 0.115 | -1.537 | 7.80601E-06 |
| 8 | Ptprc | 16.345 | 4.806 | -11.539 | 7.83063E-06 |
| 9 | Fcgr2b | 29.445 | 8.814 | -20.631 | 9.22626E-06 |
| 10 | Scimp | 4.059 | 0.557 | -3.501 | 9.33703E-06 |
| 11 | Srgn | 47.757 | 13.503 | -34.254 | 9.39187E-06 |
| 12 | Ubd | 8.176 | 0.417 | -7.760 | 9.987E-06 |
| 13 | Pik3ap1 | 16.416 | 6.053 | -10.363 | 1.10568E-05 |
| 14 | Tgm2 | 25.834 | 7.395 | -18.439 | 1.12324E-05 |
| 15 | Irf1 | 36.175 | 11.774 | -24.400 | 1.13451E-05 |
| 16 | Plac8 | 53.527 | 7.757 | -45.770 | 1.17837E-05 |
| 17 | Ccr5 | 12.634 | 4.444 | -8.190 | 1.293E-05 |
| 18 | Batf2 | 5.442 | 1.303 | -4.139 | 1.58681E-05 |
| 19 | Dusp2 | 4.600 | 1.214 | -3.386 | 1.67166E-05 |
| 20 | Fgl2 | 20.901 | 5.857 | -15.044 | 1.68507E-05 |

**Table S3.**
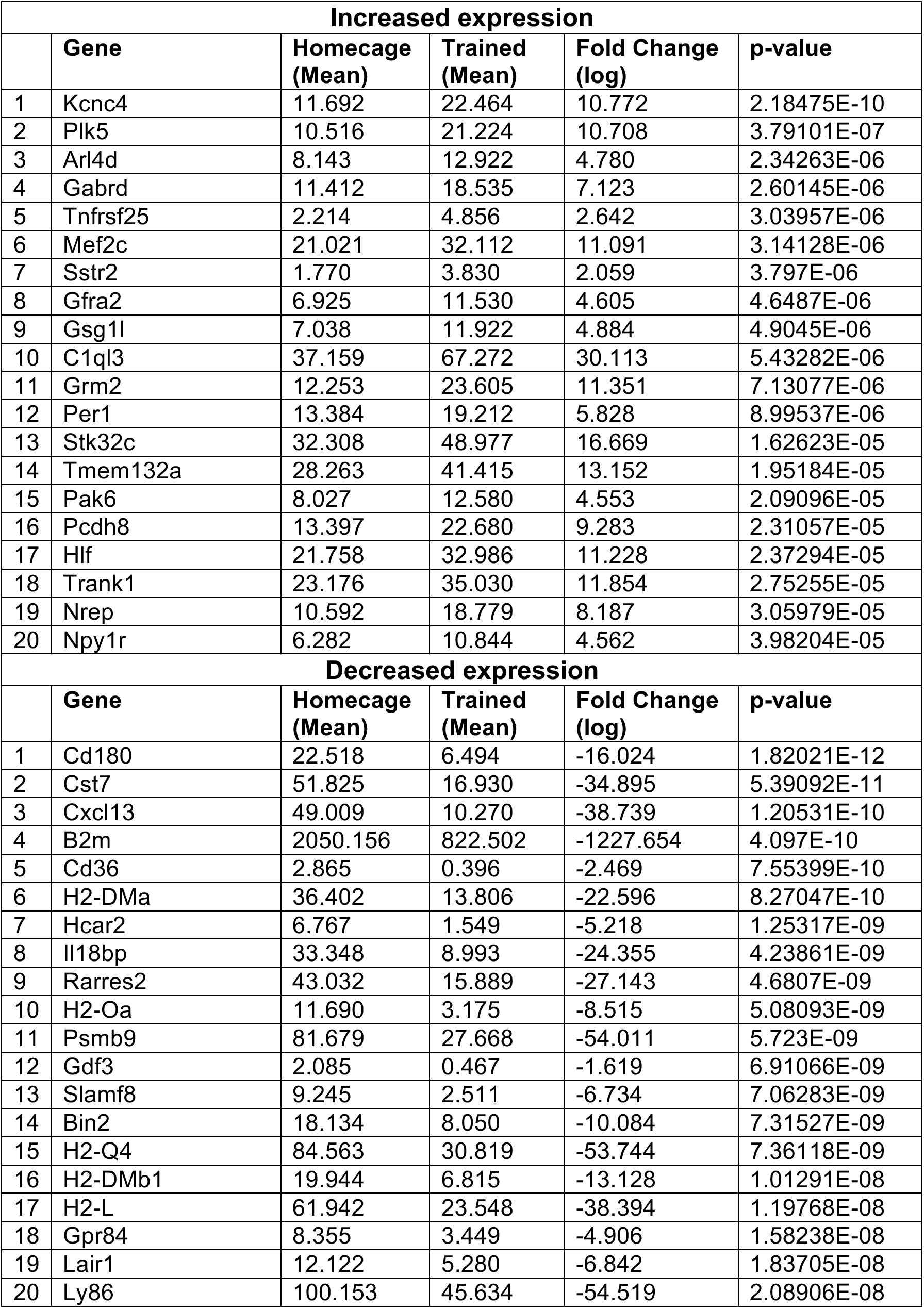
Genes with increased (top) or decreased (bottom) expression in old (18-m.o.) HDAC3^flox/flox^ mice 60m after OLM training (trained) compared to baseline (homecage). The top 20 genes increased or decreased (based on *p*-value, homecage vs trained) are shown.

**Table S4.**
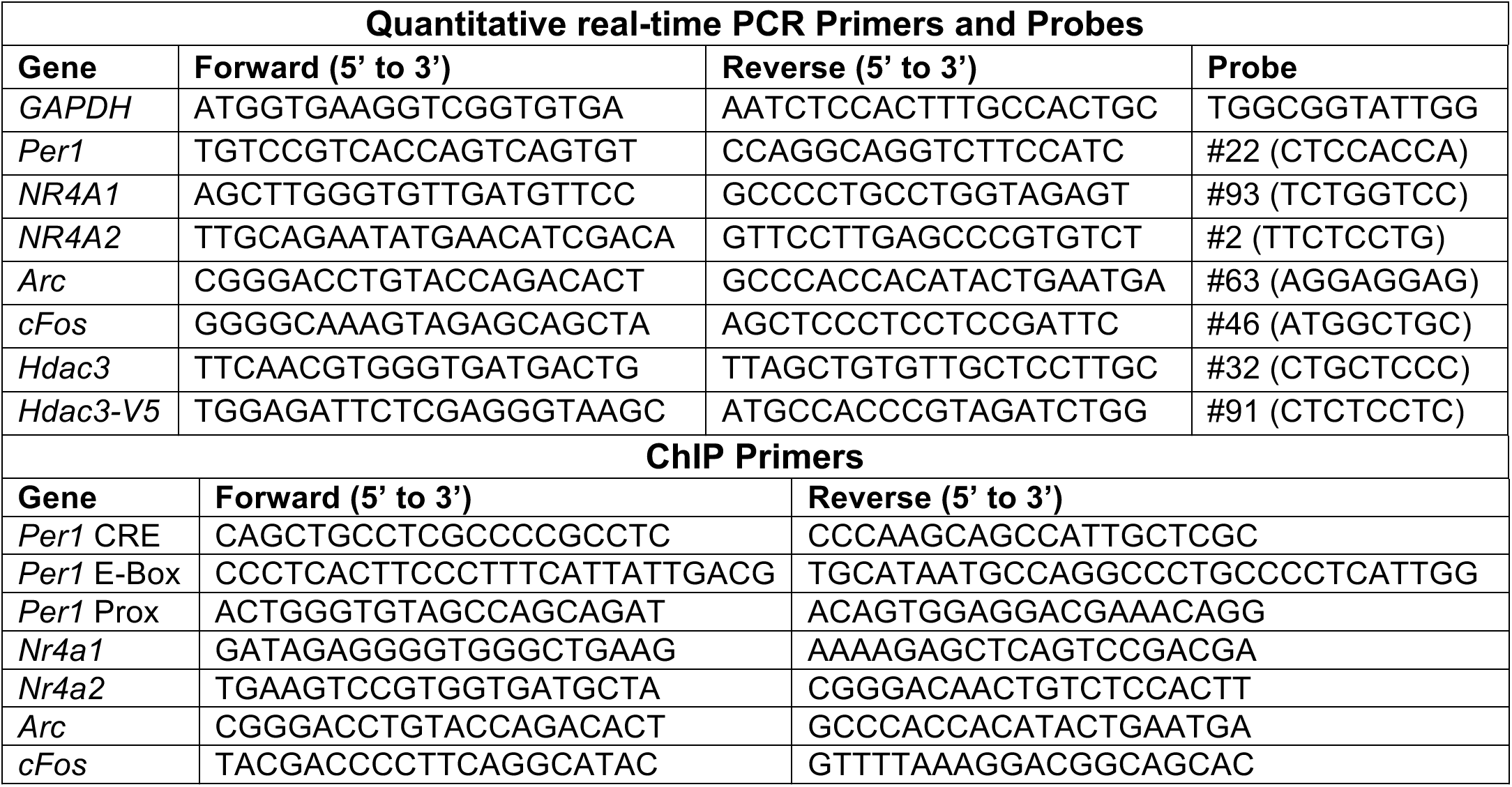
List of quantitative real-time PCR primers and probes (Roche Universal Probe Library) and ChIP primers (designed with Primer 3).

